# Fidelity of Prespacer Capture and Processing is Governed by the PAM Mediated Interaction of Cas1-2 Adaptation complex in *Escherichia coli*

**DOI:** 10.1101/602623

**Authors:** K.N.R. Yoganand, Manasasri Muralidharan, B. Anand

## Abstract

During CRISPR adaptation, short sections of invader derived DNA of defined length are specifically integrated at the leader-repeat junction as spacers by Cas1-2 integrase complex. While several variants of CRISPR systems utilise Cas4 as an indispensible nuclease for processing the PAM containing prespacers to a defined length for integration– surprisingly– a few CRISPR systems such as type I-E are bereft of Cas4. Therefore, how the prespacers show impeccable conservation for length and PAM selection in type I-E remains intriguing. In *Escherichia coli*, we show that Cas1-2/I-E– via the type I-E specific extended C-terminal tail of Cas1 –displays intrinsic affinity for PAM containing prespacers of variable length and its binding protects the prespacer boundaries of defined length from the exonuclease action that ensues the pruning of aptly sized substrates for integration. This suggests that cooperation between Cas1-2 and cellular exonucleases drives the Cas4 independent prespacer capture and processing in type I-E.

## INTRODUCTION

Prokaryotes utilize an adaptive immune response mediated by clustered regularly interspaced short palindromic repeats (CRISPR) and CRISPR associated proteins (Cas) in order to respond against infections by mobile genetic elements (MGE), *viz.*, phages and plasmids (Barrangou et al., 2007; Fineran and Charpentier, 2012; Hille et al., 2018; Horvath and Barrangou, 2010; Marraffini and Sontheimer, 2008). CRISPR encompasses a typical architecture that comprises of an array of direct repeats (∼30-40 bp), which are partitioned by short spacer sequences of viral origin. The repository of spacers acts as a vaccination card and this genetic memory acquired during pathogenic invasions guides the adaptive immune response by CRISPR-Cas (Al-Attar et al., 2011; Barrangou et al., 2007; Kunin et al., 2007). The molecular choreography of CRISPR-Cas defence is trichotomized to (I) adaptation, (II) maturation and (III) interference stages (Makarova et al., 2011). Upon phage attack, the CRISPR adaptation machinery derives short stretches of nucleic acids (prespacers) from the invaders and incorporates them into the CRISPR array. This process generates an infection memory and immunizes the host (Amitai and Sorek, 2016; Jackson et al., 2017; McGinn and Marraffini, 2019; Sternberg et al., 2016). An A-T rich leader region present upstream to the repeat-spacer array regulates the transcription of the CRISPR locus and the subsequent Cas nuclease mediated processing of this inert transcript generates short regulatory mature CRISPR RNA (crRNA) (Hochstrasser and Doudna, 2015; Punetha et al., 2018). An assemblage of crRNA with various other Cas proteins generates a surveillance complex that detects the recurring infections of MGE by base complementarity to the crRNA. This event triggers the Cas nucleases to annihilate the MGE upon repeated phage assaults and interfere with the spread of infection (Fineran and Charpentier, 2012; Horvath and Barrangou, 2010; Makarova et al., 2015; Marraffini, 2015; McGinn and Marraffini, 2019).

The CRISPR adaptation machinery expands the spacer archive, thus bestowing the host with an immunological memory to counter the ever-evolving phages. Highly conserved Cas1 and Cas2 spearhead the spacer acquisition(Jackson et al., 2017; Makarova et al., 2018; Yosef et al., 2012), however, previous investigations have also demonstrated the indispensable role of additional Cas proteins and host factors in the uptake of spacers that are derived from phage DNA and RNA (Fagerlund et al., 2017; Ivancic-Bace et al., 2015; Levy et al., 2015; Li et al., 2014b; Mohr et al., 2018; Nunez et al., 2016; Pul et al., 2010; Rollie et al., 2018; Silas et al., 2016; Wei et al., 2015b; Westra et al., 2010; Yoganand et al., 2017). Though the spacer sequences are extremely variable, the majority of the organisms (CRISPR types I, II and V) display conservation of a 2-5 nt protospacer adjacent motif (PAM) at its origin site in MGE (Heler et al., 2015; McGinn and Marraffini, 2019; Yosef et al., 2012; Yosef et al., 2013; Zetsche et al., 2015). In addition to specifying the prespacer regions for integration, PAM also guides the differentiation between self and non-self genomic regions during the interference step (Marraffini and Sontheimer, 2010). Mutations in the PAM region on MGE lead to impaired target recognition, thus evading CRISPR-Cas immune response. In such circumstances, the imperfectly paired surveillance complex at the target region directs the interference machinery and Cas1-2 to display an inflammatory immune response by rapidly acquiring more spacers by a process termed ‘primed adaptation’ (Datsenko et al., 2012; Li et al., 2014a; Li et al., 2014b; Semenova et al., 2016).

The CRISPR adaptation pathway proceeds via two sequential events: capture of the prespacer fragments from invading MGE and the site-specific integration of these captured fragments at the leader-repeat junction. The CRISPR adaptation machinery of *Escherichia coli* (type I-E) derives its spacers predominantly from the DNA debris generated during the action of multi-subunit RecBCD DNA repair complex (Levy et al., 2015). Modulation in helicase, exonuclease and endonuclease activities of RecBCD results in the production of single-stranded DNA fragments ranging from tens to thousands of nucleotides in length (Dillingham and Kowalczykowski, 2008). However, the legitimate spacers in *E. coli* are strictly 33 bp in length and abut a 5’-AAG-3’ PAM (where ‘G’ is destined to be the first residue of the spacer) (Datsenko et al., 2012; Mojica et al., 2009; Swarts et al., 2012; Yosef et al., 2012; Yosef et al., 2013). The existence of such DNA fragments is infinitesimal in the RecBCD products, hinting at the involvement of an additional processing step to generate befitting substrates. Recent studies in *Sulfolobus solfataricus* (type I-A), *Sulfolobus islandicus* (type I-A), *Bacillus halodurans* (type I-C), *Synechocystis sp*.6803 (type I-D), *Pyrococcus furiosus* (type I-G) and *Geobacter sulfurreducens* (type I-U) (Almendros et al., 2019; Kieper et al., 2018; Lee et al., 2018; Liu et al., 2017; Rollie et al., 2018; Shiimori et al., 2018; Zhang et al., 2019) highlighted the indispensable role of Cas4 nuclease in PAM selection and prespacer processing. The occurrence of Cas4 is predominantly limited to type I CRISPR-Cas system with the exception of subtypes I-E and I-F (Makarova et al., 2018). In one case, it was observed that an extended variant of Cas2 with an unorthodox C-terminus DnaQ exonuclease domain assists *Streptococcus thermophilus* DGCC7710 (type I-E) in prespacer trimming (Drabavicius et al., 2018). Cas2-DnaQ domain fusion is non-ubiquitous and even model organisms for spacer acquisition studies like *E. coli* (type I-E) do not harbour it. Though recent studies envisage the involvement of exonucleases during spacer acquisition in *E. coli* (Radovcic et al., 2018), the molecular events guiding PAM selectivity and prespacer processing remain obscure.

Upon production of legitimate prespacers, Cas1-2 integrase complex catalyses the prespacer incorporation at the leader adjoining repeat (Li et al., 2014b; McGinn and Marraffini, 2016; Nunez et al., 2015a; Nunez et al., 2015b; Rollie et al., 2015; Swarts et al., 2012; Wei et al., 2015b; Yosef et al., 2012). This polarized mechanism records the chronology of infections by positioning the recent spacer towards a promoter encompassing leader. During recurring infections, swift expression of the latest spacers ensures a productive fight by a rapid and robust immune response (McGinn and Marraffini, 2016). In conjunction with Cas1-2, various conserved DNA motifs present in the leader and repeat regions mandate the fidelity of prespacer integration (Arslan et al., 2014; Goren et al., 2016; McGinn and Marraffini, 2016; Moch et al., 2017; Nunez et al., 2016; Wang et al., 2016; Wei et al., 2015a; Yoganand et al., 2017; Yosef et al., 2013). In *E. coli* (type I-E), upon interaction with these conserved regions, a sequence-specific genome architectural protein termed integration host factor (IHF) restructures the leader and generates a docking site for Cas1-2 integrase at the leader-repeat junction (Nunez et al., 2016; Wright et al., 2017; Yoganand et al., 2017). In contrast, the presence of Cas1-2 binding site at the leader proximal region obviates the involvement of host factors during spacer acquisition in CRISPR type II-A system (McGinn and Marraffini, 2016; Wright and Doudna, 2016; Xiao et al., 2017).

Unlike many of the type I CRISPR-Cas encompassing prokaryotes, *E. coli* (type I-E) lacks Cas nucleases like Cas4 and Cas2-DnaQ to generate processed prespacers for efficient homing at CRISPR locus (Makarova et al., 2018). Nevertheless, this inadequacy doesn’t appear to hinder PAM selection or spacer size preference (Mojica et al., 2009; Shipman et al., 2016; Swarts et al., 2012; Yosef et al., 2012; Yosef et al., 2013). Intrigued by these observations, we sought to understand how prespacers are selected and tailored to the appropriate size for CRISPR adaptation. Here, we demonstrate that the PAM directed interactions with longer DNA fragments signals Cas1-2 to demarcate potential prespacer boundaries. Upon supplementing the reaction with exonucleases to mimic the cellular environment, we found that Cas1-2-DNA nucleoprotein complex could protect DNA fragments of ∼33 bp length. Further, we show that these protected fragments could be efficiently integrated into the CRISPR locus. These findings demystify the mechanism by which *E. coli* efficiently scales the fragments of the foreign DNA to generate viable prespacers of desired length, in contrast to other CRISPR-Cas subtypes that possess dedicated prespacer processing nucleases such as Cas4.

## RESULTS

### Length of the prespacers dictates their integration at the CRISPR array

We utilised previously established *in vitro* spacer integration assay (Yoganand et al., 2017) to understand the prespacer parameters that necessitate their uptake during CRISPR adaptation. In this assay, we employed Cas1-2 integrase, IHF, prespacers and linear CRISPR DNA (70 bp leader followed by two repeat-spacer units) (Figure 1A and S1A). Generally, prespacer integration proceeds via a trans-esterification reaction, wherein 3’-OH of the prespacer makes a nucleophilic attack at the target site to get itself ligated (Nunez et al., 2015b; Rollie et al., 2015). Upon Proteinase-K treatment to release CRISPR DNA from Cas1-2 and IHF interactions, we monitored the outcome of this ligation by electrophoretic mobility shift assays (EMSA) (Figure 1C and S1B).

**Figure 1.**
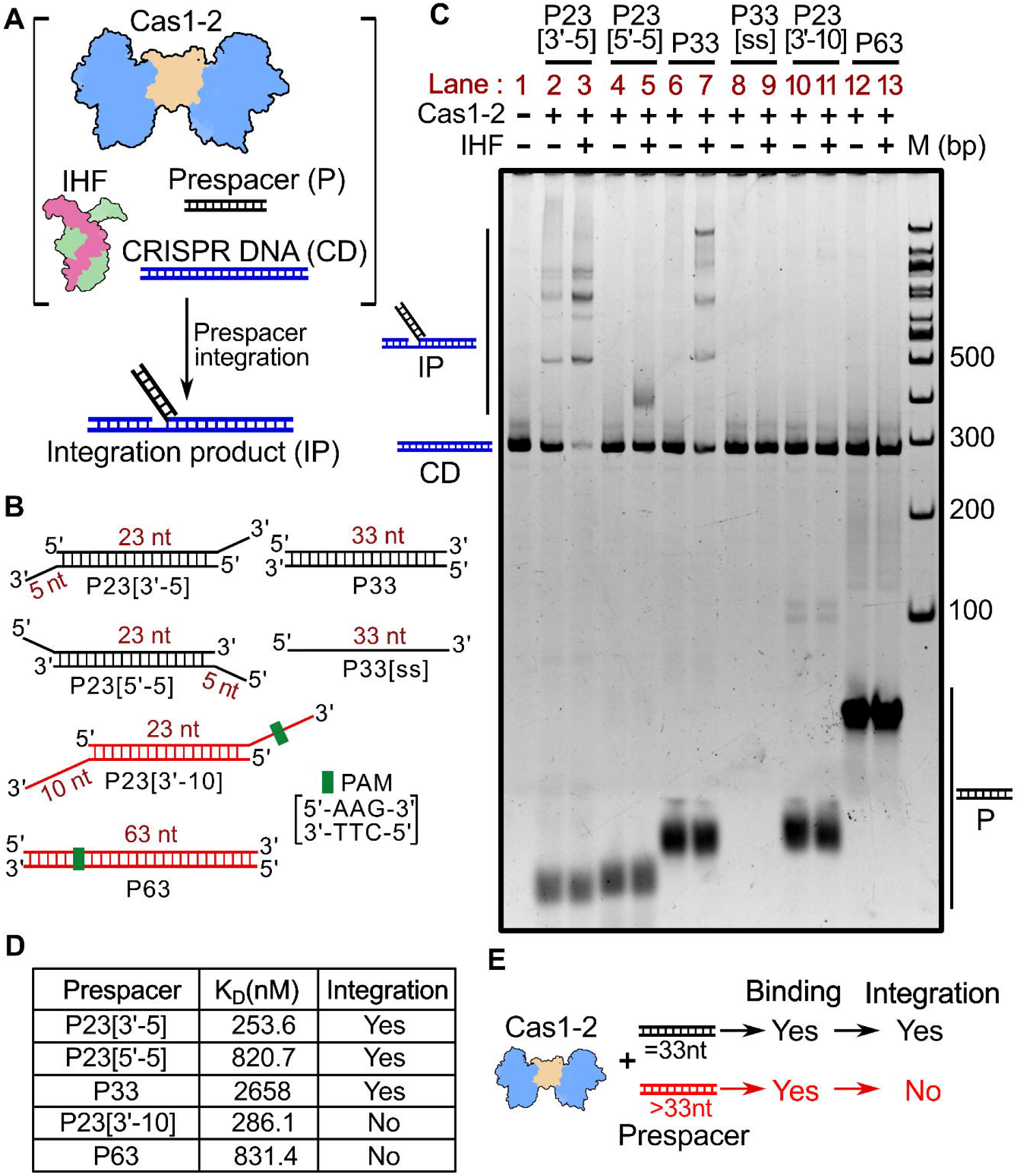
Prespacer length regulates the fate of spacer integration at CRISPR locus. (A) Schema of Cas1-2 mediated prespacer integration assay is presented. Integration product (IP) and various reaction components such as Cas1-2 (in blue and brown), IHF (in pink and green), prespacer (Black ladder), CRISPR DNA (Blue ladder) are pictorially depicted. (B) Duplex (P33 and P63), partial duplex (3’-overhang: P23[3’-5] & P23[3’-10]; 5’-overhang: P23[5’-5]) and single stranded (P33[ss]) prespacers that are employed in the integration assay are shown. Prespacers with overall length of 33 nt are coloured black whereas those with >33 nt length are displayed in red colour. Position of PAM is shown as green box. (C) Native gel displaying the proteinase K treated samples of spacer integration assay is shown. Prespacers P23[3’-5] (lanes 2-3), P23[5’-5] (lanes 4-5), P33 (lanes 6-7), P33[ss] (lanes 8-9), P23[3’-10] (lanes 10-11) and P63 (lanes 12-13) were employed in integration reactions. Absence (–) or presence (+) of Cas1-2 and IHF is indicated on top of each lane. Positions of bands corresponding to CRISPR DNA (CD), integration product (IP) and prespacer (P) are pictorially represented. Whereas DNA molecular weight marker (M) positions are shown on the right side. (D)The equilibrium disassociation constant values (K_D_) of Cas1-2 with each type of prespacer substrate are displayed. The success (Yes) or failure (No) of integration for each prespacer is shown. (E) Cartoon depicting the relationship between protospacer length (33 nt and >33 nt) and Cas1-2 (in blue and brown) mediated binding and integration is presented. Possibility of binding and integration of each prespacer is denoted as ‘Yes’ or ‘No’.

Spacers in *E. coli* are routinely derived from the remnant DNA fragments generated post-RecBCD mediated double-stranded break repair (Levy et al., 2015). This process results in sheared ssDNA fragments that can range up to kilobases in length (Dillingham and Kowalczykowski, 2008). Upon reannealing to their complementary sequences, various sizes and forms of DNA could be generated that encompass blunt ends, 3’ or 5’ overhangs. Previous *in vivo* studies also demonstrated the incorporation of 33 bp spacer regions derived from longer electroporated DNA fragments of 63 bp in length (Shipman et al., 2016). To simulate these conditions *in vitro*, we employed various types of DNA fragments such as P33 (33 bp duplex), P33[ss] (33 nt ssDNA), P23[3’-5] (23 bp duplex with 5 nt 3’ overhangs), P23[5’-5] (23 bp duplex with 5 nt 5’ overhangs), P23[3’-10] (23 bp duplex with 10 nt 3’ overhangs) and P63 (63 bp blunt duplex) in spacer integration assays (Figure 1B). Upon incubation with Cas1-2 and IHF, we could identify slow migrating integration products that appear to be larger than the CRISPR DNA in each of the reactions that contain P23[3’-5] or P23[5’-5] or P33 (Lanes 3, 5 and 7 in Figure 1C). Strikingly, bands corresponding to integration products weren’t found when we substituted the reaction mixtures with P33[ss] or P23[3’-10] or P63 (Lanes 9, 11 and 13 in Figure 1C). These findings suggest that either duplex (P33) or partial duplex (comprising of 3’ overhang (P23[3’-5]) or 5’ overhang (P23[5’-5])) prespacers with an effective length of 33 nt are strictly required during CRISPR adaptation. This bias in prespacer size preference could possibly arise due to the weakening of Cas1-2 interaction with long substrate precursors (such as P23[3’-10] and P63 with effective length of 43 nt and 63 nt, respectively) and/or inefficient integration of such DNA fragments at the target site in CRISPR locus. Previous structural studies showed that Cas1-2 could interact with 33 nt partial duplex prespacers. Here, EMSA performed with Cas1-2 integrase and various prespacers (Figure S2 and 1D) showed that P23[3’-10] (K_D_ = 286.1 nM) displays comparable affinity to P23[3’-5] (K_D_ = 253.6 nM). In contrast to this, the affinity of P63 (K_D_ = 831.4 nM) for Cas1-2 is stronger than that of P33 (K_D_ = 2.658 μM). These experiments clearly suggest that Cas1-2 can interact with DNA fragments of varied lengths; however, the integration at the target site could be achieved only in the presence of DNA fragments with an effective length of 33 nt (Figure 1E).

### Cas1-2 foothold protects potential prespacer regions during exonuclease action

CRISPR adaptation in *E. coli* mandates precisely sized prespacers (Figure 1). To cater to this need, long DNA fragments generated during RecBCD repair have to be further trimmed by nuclease action. Of the multitude of proteins encoded by the Cas operon, only Cas1 and Cas2 contribute to naïve spacer acquisition in *E. coli*. We could not notice processing of longer prespacers (P23[3’-10] and P63) even at higher Cas1-2 concentrations (Figure S3). The type I-E system is devoid of prespacer processing exonuclease Cas4 and Cas1-2 by itself could not complement the absence of Cas4 (Figure S3). Hence, we predicted the involvement of cytoplasmic nucleases in trimming the longer prespacer to a suitable length. Having established the fact that Cas1-2 indeed binds the DNA fragments of variable length (Figure 1D and S2), we sought to test the fate of Cas1-2 bound DNA fragments upon supplementing exonucleases. To achieve this, we allowed the binding of Cas1-2 with a longer prespacer P63 and then treated this nucleoprotein complex with a mixture containing 5’→3’ acting T5 exonuclease (T5exo) and 3’→5’ acting Exonuclease III (ExoIII) (Figure 2A). To our surprise, we identified a smear of protected DNA fragments (P63exo+) that ranged from 30 to 40 nt in the sample containing both P63 and Cas1-2 (Lane 8 in Figure 2A). Whereas such protection could not be seen when we treated P63 in the absence of Cas1-2 or in presence of Cas1 or Cas2 alone (Lanes 5, 6 and 7 in Figure 2A). Coincidentally, the length of the protected fragments corresponded to legitimate spacer size in *E. coli* (∼33 nt). We wondered whether these trimmed DNA fragments could act as potential prespacers for integration into the CRISPR array. To test this presumption, we purified and utilised P63exo+ DNA fragments as prespacers in spacer integration assay. In line with previous experiment (Figure 1A), we could not observe any integration events when we employed longer prespacer P63 (Lane 7 in Figure 2B). Surprisingly, in case of purified P63exo+ fragments, we identified slow migrating integrated products (Lane 9 in Figure 2B). By recapitulating these observations, we could suggest that Cas1-2 mediated binding of large DNA fragments secure the boundaries of suitable prespacers from the exonucleolytic action of cellular nucleases in *E. coli* (Figure 2C).

**Figure 2.**
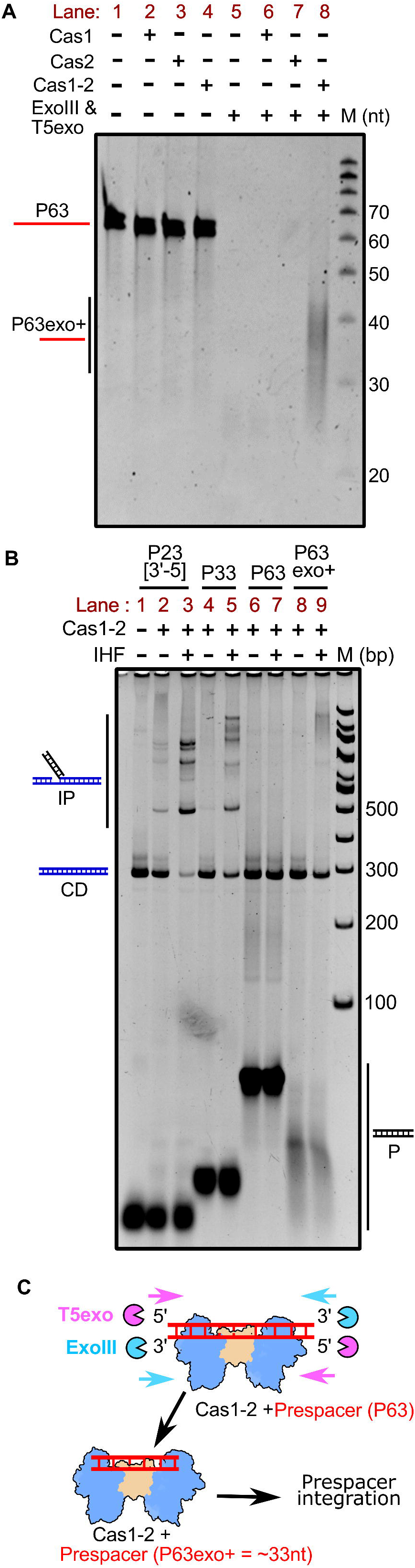
Tailoring of Cas1-2 bound large DNA fragments by exonucleases generates integration competent prespacers. (A) Denaturation gel depicting the nuclease treatment of Cas1-2 bound P63 DNA fragments is displayed. Presence (+) or absence (-) of each reaction component is labelled on top of each lane. Positions corresponding to the substrate (P63) and T5 exo/ExoIII digested DNA fragments (P63exo+) are indicated on the left side, whereas oligo marker (M) positions are shown on the right. (B) Native gel displaying the integration reactions employing various prespacers (P23[3’-5] (lanes 1-2), P33 (lanes 3-4), P63 (lanes 6-7)) and Cas1-2 protected DNA fragments (P63exo+) (lanes 8-9) is shown. Presence (+) or absence (-) of each reaction component is labelled on top of each lane. Positions corresponding to CRISPR DNA (CD) and integrated products (IP) are pictorially depicted on the left, whereas prespacer (P) and DNA molecular weight marker (M) positions are shown on the right. (C) Schema illustrating the mechanism of Cas1-2 mediated protection of prespacer boundaries is displayed. Cas1-2 (in blue and brown), T5 exonuclease (magenta pie), Exonuclease III (Cyan pie) and prespacer P63 (red ladder) are pictorially portrayed.

### PAM directed binding of Cas1-2 defines the boundary for prespacers

Since Cas1-2 binding was shown to mark the spacer boundaries (Figure 2), we sought to identify and map these protected regions. To accomplish this, we utilised variants of 63 bp blunt-ended prespacers that encompass fluorescein labelled 3’-end (6-FAM) either on the top (P63T*) or on the bottom (P63B*) strand (Figure 3A). Further, in order to identify the footprints of Cas1-2 on these 3’-end labelled prespacers, we incubated Cas1-2 bound prespacer complex with T5exo. Here, the Cas1-2 binding on the prespacer acts as a roadblock and stalls the 5’→3’ progression of T5exo. The length of the resultant labelled fragments specifies the stalling points of the exonuclease, which in turn, indicates the binding position of Cas1-2 on prespacer. Utilizing this approach, we mapped the cleavage termination points on the top and bottom strands of the prespacer. After T5exo treatment of P63T*, we could observe ∼28 nt labelled fragment. This is indicative of an inherent nuclease stalling point (in absence of roadblocks such as Cas1-2) around the 28^th^ nt position from the labelled end in P63T* (Lane 2 in Figure 3B). However, owing to complete exonucleolytic cleavage of the bottom strand we didn’t observe such stalling on P63B* (Lane 10 in Figure 3B). Upon T5exo treatment of Cas1-2 bound P63T* complex, we noticed a shift in the nuclease stalling point to ∼45 nt position from the labelled end (Lane 4 in Figure 3B). This finding maps the Cas1-2 binding position to be around 45 nt from the labelled end (P63T* in Figure 3B). Coincidentally, this binding position of Cas1-2 on P63T* is localized around a cognate PAM sequence (5’-AAG-3’ ranging from 47 nt to 49 nt upstream of labelled position) (P63T* in Figure 3A). Prompted by this finding, we were interested to identify the extent of protection that Cas1-2 could confer upon its binding at PAM region. To accomplish this, we treated the Cas1-2 bound P63B* complex with T5exo. Here, the resulting length of the protected fragments upon exonuclease treatment indicated that Cas1-2 complex interaction can guard a region spanning ∼45 nt from the labelled end in P63B* (Lane 12 in Figure 3B and P63B* in Figure 3A). As the PAM residues are positioned at 14 nt from the labelled end of the P63B*, the effective length of the protected prespacer from the PAM is ∼30 nt. Overall these findings demonstrate a probable mechanism by which Cas1-2 could selectively acquire 33 nt prespacers that are bordered by the PAM as in *E. coli*.

**Figure 3.**
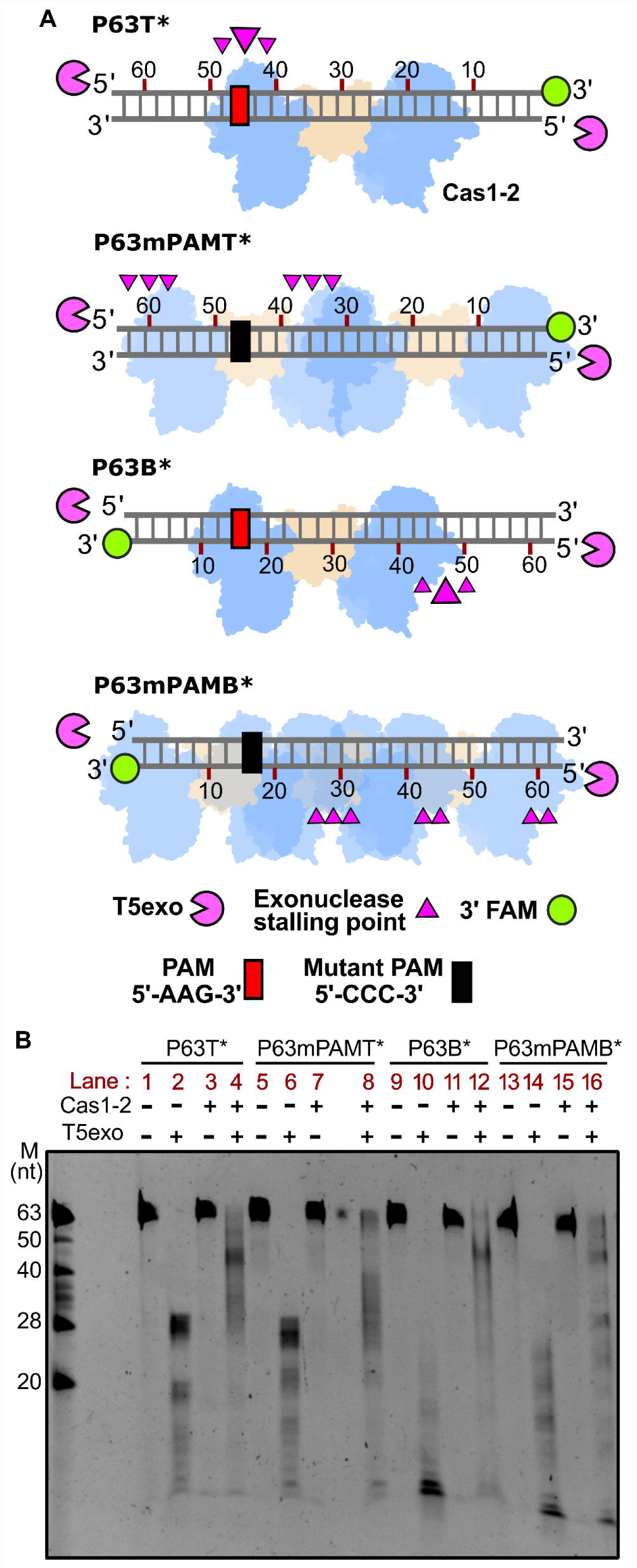
Cas1-2 complexes are predominantly localized around PAM region. (A) Schematic representation depicting various fluorescein labelled prespacer substrates (P63T*, P63mPAMT*, P63B* and P63mPAMB* as grey ladder) used in the assay is shown. Labelled 3’-end of each prespacer is highlighted as green circles, whereas PAM and its mutated positions are depicted as red and black boxes, respectively. Numbering on the DNA represents the distance (in nt) of particular position from the labelled end. T5 exo (magenta pie) is positioned at susceptible 5’-ends of DNA substrate. Positions of T5exo stalling points (magenta triangles) and binding sites of Cas1-2 (blue and brown blobs) that are estimated from nuclease footprinting assay performed in (B) are pictorially displayed. (B) Denaturation gel depicting the T5exo treatment of Wt. Cas1-2 bound fluorescein labelled P63 variants (P63T*, P63mPAMT*, P63B* and P63mPAMB*) are displayed. Presence (+) or absence (-) of each reaction component is labelled on top of each lane. Positions corresponding to the DNA fragments of oligo marker (M) are shown on the left.

To reinforce these observations, we employed mutated P63 DNA fragments (P63mPAMT* and P63mPAMB* in Figure 3A) that are devoid of any cognate *E. coli* PAM sequence (5’-AWG-3’, where W=A/T). These labelled fragments were incubated with Cas1-2 and later treated with T5exo. Here, we found large smears and multiple bands upon employing P63mPAMT* (Figure 3A and Lane 8 in 3B) and P63mPAMB* (Figure 3A and Lane 16 in 3B). The varied length of resultant labelled fragments is indicative of numerous stalling points on these P63 mutants (Figure 3A) that occurred due to Cas1-2 binding.

These results highlight that the PAM plays a key role in defining the prespacer boundaries. When a larger DNA fragment encompasses a PAM sequence, Cas1-2 mediated binding and subsequent nuclease action generates prespacers that are particularly bordered by PAM (Lanes 4 and 12 in Figure 3B; P63T* and P63B* in Figure 3A). Whereas such specificity is lost when the DNA fragments lack PAM. Here Cas1-2 seems to interact with various regions of the DNA and result in the generation of illicit prespacers that do not encompass PAM at the desired position (Lanes 8 and 16 in Figure 3B; P63mPAMT* and P63mPAMB* in Figure 3A).

### Intrinsic specificity of Cas1-2 circumvents the requirement of Cas4 during PAM selection in *E. coli*

Having found the role of PAM mediated interaction in selecting the prespacers for uptake, we attempted to understand the intrinsic molecular principles that confer precision to Cas1-2 in PAM selectivity and prespacer scaling. Previous structural studies of CRISPR adaptation complex in *E. coli* suggested that the extended Cas1 C-terminal tail of apoCas1-2 complex (Nuñez et al., 2014) gets organized around the PAM residues upon binding to prespacer DNA (Figure S4A and S4B; (Wang et al., 2015). In particular, Q287 and I291 residues of this proline-rich C-terminal tail make direct contacts with the nucleotides of PAM (Figure S4B), thus possibly imparting the PAM specificity. In another striking feature, a pair of Y22 residues that are derived from two different Cas1 protomers scales a 23 bp duplex region of prespacer by stacking interactions at either ends (Figure S4C) (Nunez et al., 2015a; Wang et al., 2015). This gating mechanism at the Cas1-2 platform seems to adjudge the spacer length by facilitating the positioning of 3’-overhang at the catalytic groove for integration (Figure S4C). To validate these presumptions, we employed Cas1-2 variants that encompass either deletion of a Cas1 C-terminal tail (ΔC - ΔP279-S305) or a Cas1 with alanine substitution at tyrosine 22 (Y22A) (Figure S5A) and analysed their footprints upon binding to P63B* and P63T* (Figure 4A and S6E). As a control, we also used a Cas1-2 variant (Cas1 Penta-mutant (5M - Q24H, P202Q, G241D, E276D, L297Q)) (Figure S4D) that was previously shown to abrogate PAM selectivity (Shipman et al., 2016).

**Figure 4.**
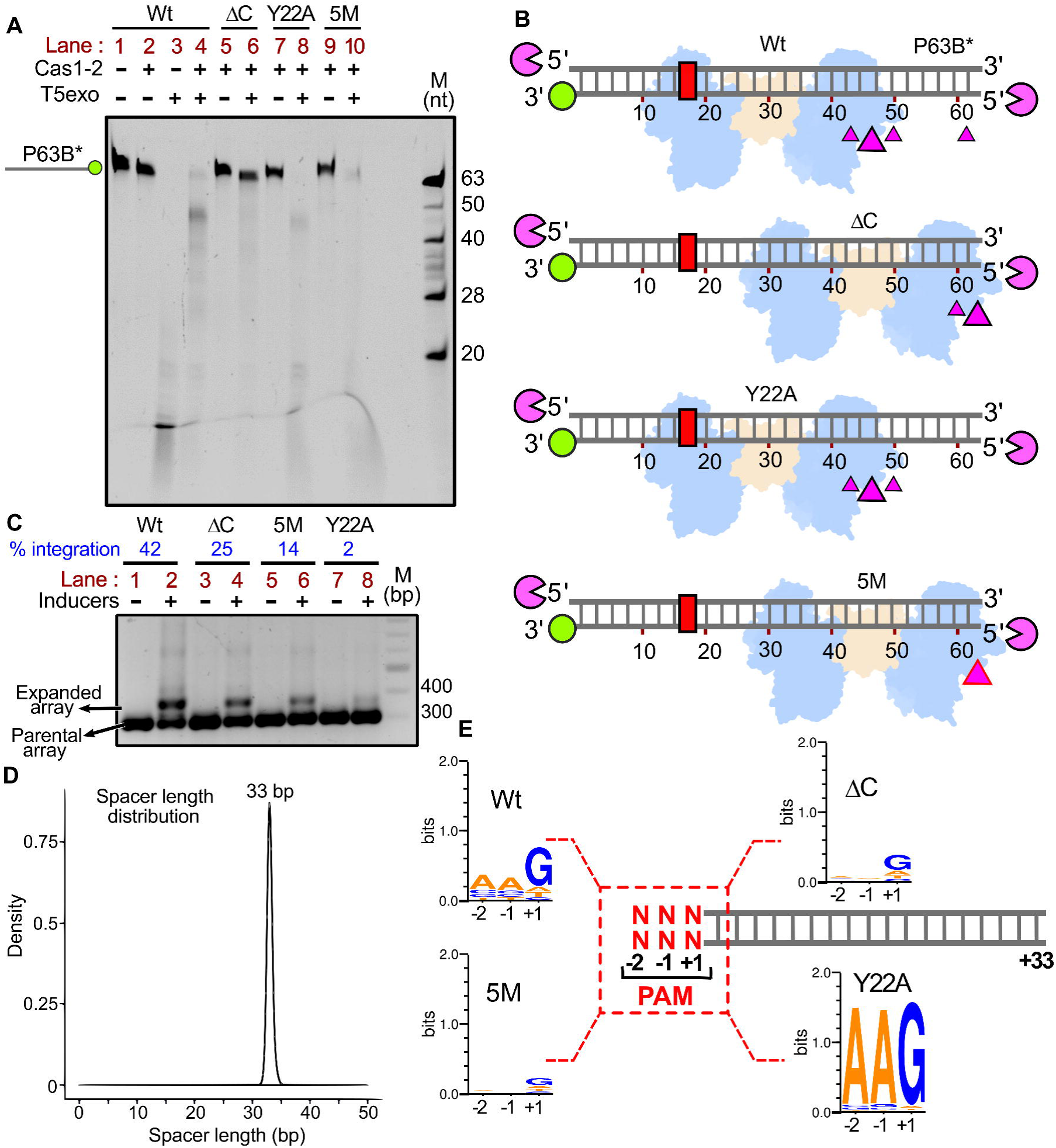
Intrinsic specificity of Cas1-2 integrase direct the uniformity in spacer length and PAM preference during CRISPR adaptation. (A) Denaturation gel depicting the T5exo treatment of Cas1-2 (Wt (lanes 1-4) or ΔC (lanes 5-6) or Y22A (lanes 7-8) or 5M (lanes 9-10)) bound fluorescein labelled P63B* is displayed. Presence (+) or absence (-) of each reaction component is indicated on top of each lane. Position of P63B* labelled DNA fragment is shown on the left, whereas oligo marker (M) positions are indicated on the right. (B) Schematic illustration depicting the footprinting assay performed in (A) is displayed. DNA substrate P63B* (grey ladder), positions of 3’ fluorescein label (green circle) and PAM region (red rectangle) are pictorially represented. Numbering on the DNA represents the distance (in nt) of particular position from the labelled end. T5 exo (magenta pie) is positioned at susceptible 5’-ends of DNA substrate. Positions of T5exo stalling points (magenta triangles) and binding sites of each variant of Cas1-2 (Wt or ΔC or Y22A or 5M in blue and brown blobs) that are estimated from nuclease footprinting assay performed in (A) are pictorially displayed. (C) Agarose gel depicting the PCR products from spacer acquisition assay performed in *E. coli* harbouring the plasmids that express the Cas1-2 variants (Wt (lanes 1–2), ΔC (lanes 3–4), 5M (lanes 5–6) and Y22A (lanes 7-8)) is shown. Absence (–) or presence (+) of inducers is indicated on top of each lane. Positions corresponding to parental and expanded arrays (CRISPR 2.1 array) are indicated on left. The extent of integration ([(Amount of expanded array) / (Amount of parental array + Amount of expanded array)] * 100) is displayed on top of the respective lanes (indicated in blue) whereas DNA marker (M) positions are represented on the right. (D) Overlay of plots depicting length distribution of the newly acquired spacers that are incorporated into CRISPR 2.1 array by Cas1-2 variants (Wt, ΔC, 5M and Y22A) during *in vivo* integration assay (C) is shown. X-axis depicts length of the spacer (nt) whereas normalised frequency (density) is indicated on the Y-axis. (E) Illustration depicting the PAM preference of Cas1-2 variants (Wt, ΔC, 5M and Y22A) during *in vivo* integration assay (C) is displayed. +1 to +33 sequence of each spacer (grey ladder) was extracted from high-throughput sequencing data. Subsequently, sequence information of −1 and −2 positions of each spacer was derived from the respective plasmid/genome sequence. The conservation profile of PAM sequences (−2, −1 and +1 in red) corresponding to the respective Cas1-2 variant is shown as a sequence logo.

Similar to Wild-type Cas1-2 (Wt), T5exo treatment of Y22A bound P63B* led to the generation of a nuclease stalling point at ∼45 nt albeit with reduced efficiency (Compare lanes 4 and 8 in Figure 4A). These findings indicate that the alanine replacement of Cas1 Y22 residue does not result in the altered PAM specificity (Wt and Y22A in Figure 4B). However, the affinity of Y22A for the P63 is altered (Y22A K_D_ = 6.22 μM vs Wt K_D_ = 831.4 nM) (Figure S6C and S6D), and this suggests that the absence of Y22 mediated stacking interaction seems to reduce its prespacer protection ability against the nuclease action. Unlike Wt, nuclease stalling points were observed around 60 nt from the labelled ends of P63B* upon employing ΔC and 5M Cas1-2 variants (Lanes 6 and 10 in Figure 4A). Owing to its low affinity, 5M seems to display reduced protection of prespacers from nuclease action (ΔC K_D_ = 7.584 μM vs 5M K_D_ = 21.59 μM) (Figure S6A and S6B). The shift in this nuclease stalling point indeed indicates that the ΔC and 5M variants display an impaired PAM specificity and were randomly interacting at the ends of P63B* (ΔC and 5M in Figure 4B). To fortify our observations, we sought to understand the impact of these mutations on the composition of acquired spacers *in vivo*. To achieve this, we induced spacer acquisition in *E. coli* IYB5101 by expressing Cas1-2 variants. Here, we observed that the mutations in ΔC and 5M have partly reduced the spacer incorporation efficacy of Cas1-2 (Compare Lanes 2, 4 and 6 in Figure 4C). Surprisingly, Y22A displayed a drastic reduction of spacer uptake *in vivo* (Compare Lanes 2 and 8 in Figure 4C), despite the fact that this mutation could not effectively avert spacer integration *in vitro* (Figure S5B). Expanded CRISPR arrays corresponding to the expression of each mutant were purified and the sequences of newly incorporated spacers were derived from high-throughput sequencing. In line with previous studies, we observed that the spacers were originated from both genome and plasmid (Mojica et al., 2009; Shipman et al., 2016). Irrespective of these mutations in Cas1-2, the length of the incorporated spacers is strictly conserved (i.e., 33 nt) (Figure 4D). This finding suggests that Y22 mediated stacking interaction with prespacer or the Cas1 C-terminal restructuring is dispensable for the scaling of prespacers. These spacer sequences were matched with that of the plasmid and genome to identify the adjoining PAM nucleotide sequences. Despite the display of precise prespacer scaling by Cas1-2 variants, the specificity towards PAM region appears to be highly altered (Figure 4E). In concurrence with previous studies, we observed that most of the spacers acquired by Wt Cas1-2 encompass a conserved PAM region (5’-AAG-3’, where ‘G’ indicates +1 position of 33 nt spacer) (Figure 4E). In line with the nuclease protection assay (Fig 4A and 4B), we didn’t observe any preference towards the PAM region when we employed 5M or ΔC (Figure 4E). This finding bolsters the involvement of Cas1 C-terminal tail in PAM selectivity. Despite the reduced efficiency in spacer acquisition *in vivo* (Figure 4C), surprisingly, Y22A has displayed a striking precision for PAM selectivity, suggesting that this mutation bestowed high fidelity with respect to PAM recognition (Figure 4E).

## DISCUSSION

CRISPR system in *E. coli* (type I-E) displays precise scaling of prespacer length and stringent selectivity of PAM (Shipman et al., 2016; Swarts et al., 2012; Yosef et al., 2013). Here, we attempted to uncover the elusive molecular events that drive the generation of competent substrates for homing at the leader-repeat junction by CRISPR adaptation complex. During naïve adaptation, prespacers are predominantly foraged from the DNA fragments generated during RecBCD mediated repair (Levy et al., 2015). Whereas Cas3 mediated DNA resection appears to surrogate the RecBCD role during primed adaptation (Datsenko et al., 2012; Künne et al., 2016; Semenova et al., 2016). Being helicase-nuclease enzymes, RecBCD and Cas3 result in a varied length of ssDNA fragments (Dillingham and Kowalczykowski, 2008; Sinkunas et al., 2011). Despite the fact that ssDNA fragments were found to be associated with Cas1 *in vivo* (Musharova et al., 2017), their integration at the CRISPR locus has been highly inefficient in comparison to duplex and partial-duplex DNA forms (Figure 1) (Nunez et al., 2015b). This points to a certitude that duplex formation by annealing of complementary ssDNA strands upon helicase-nuclease action could potentially generate prespacers that elicits CRISPR adaptation. Here, we demonstrate that irrespective of being a duplex or partial duplex the overall length of the prespacer must be of the spacer size (i.e., ∼33 nt) for a successful integration (Figure 1). Though being incompetent for integration *in vitro* (Figure 1B), 63 bp long DNA fragments electroporated into *E. coli* that harbours Cas1-2 led to the acquisition of 33 bp spacers and these were found to be derived from the 63 bp long oligonucleotides (Shipman et al., 2016). Unlike the precise DNA trimming displayed *in vivo*, Cas1-2/I-E alone is inept of such activity upon incubation with longer DNA fragments (P63 in Figure S3). In contrast to previous observation (Wang et al., 2015), despite the presence of PAM in 10 nt 3’ overhang of P23[3’-10], we could not detect prespacer processing even at elevated Cas1-2 concentrations (P23[3’-10] in Figure S3). Similar to *E. coli* (type I-E), when Cas1-2 of *B. halodurans* (type I-C) and *S. thermophilus* DGCC7710 (type I-E) is presented with prespacers of legitimate length, the requirement of Cas4 and DnaQ, respectively, is obviated and successful integration is observed (Drabavicius et al., 2018; Lee et al., 2018). These observations fortify the fact that the Cas1-2 alone catalyse the spacer integration, whereas nuclease action of Cas4 or DnaQ supplement this process by ensuring the generation of prespacers with a compatible length for integration. Notably, Cas1 and Cas2 are the only proteins of the *cas* operon that are indispensable for naïve adaptation in *E.* coli (Datsenko et al., 2012; Yosef et al., 2012). Since they are incapable of trimming the prespacer to the desired length (Figure S3), Cas1 and Cas2 alone can’t complement the dearth of exclusive prespacer processing Cas nucleases (*viz*., Cas4 and an extended variant of Cas2 with auxiliary DnaQ domain fusion). These observations emphasize the involvement of cellular nuclease(s) in the generation of efficient prespacers for integration. Structures of *E. coli* Cas integrase complex reveals that ∼33 bp length of DNA can be exactly accommodated in between two active site regions of Cas1-2 (Nunez et al., 2015a; Wang et al., 2015). This hints at the fact that the Cas1-2 foothold can only mask 33 bp region and the remainder of bound DNA fragments is exposed to potential nuclease action. Mimicking such conditions, here we incubated Cas1-2 bound longer DNA fragments (P63) with 3’→5’ (ExoIII) and 5’→3’ (T5exo) acting exonucleases. These reactions resulted in the generation of fragments that are in the range of cognate *E. coli* spacer size (Figure 2A). Furthermore, these processed DNA fragments were readily utilised as substrates by Cas1-2 integrase (Figure 2B). This implies that the lack of bona fide prespacer processing Cas4 in type I-E system of *E. coli* appears to be complemented by the action of cellular nucleases. Owing to the presence of a plethora of nucleases in *E. coli*, additional investigations are necessary to identify if a single nuclease or a conglomerate action of several nucleases could lead to the generation of competent prespacers for CRISPR adaptation.

In most of the prokaryotes, specificity of CRISPR adaptation machinery drives the uptake of phage-origin prespacers that are bordered by a PAM sequence (McGinn and Marraffini, 2019). Though the removal of Cas4 in *S. islandicus* (type I-A), S*ynechocystis sp*.6803 (type I-D) and *P. furiosus* (type I-G) didn’t hamper spacer uptake completely, deficiency of this nuclease led to plummeted PAM preference (Kieper et al., 2018; Shiimori et al., 2018; Zhang et al., 2019). Likewise, mutations in metal coordinating residues (K102A and D87A) of RecB domain of Cas4-Cas1 fusion from *G. sulfurreducens* (type I-U) led to skewed PAM specificity (Almendros et al., 2019). Moreover, *in vitro* studies performed with adaptation complex of *B. halodurans* (type I-C) and *S. solfataricus* (type I-A) had revealed that Cas4 nuclease avoids the processing of free DNA ends that are devoid of PAM sequence (Lee et al., 2018; Rollie et al., 2018). This preferential activity of Cas4 seems to act as a critical checkpoint in ensuring the productive uptake of infection memory by Cas1-2 in the hosts. Unlike in type I and III, prokaryotes containing type-II CRISPR-Cas system wield a fewer number of Cas proteins (*viz.*, Cas1, Cas2, Cas9 and Csn2) to mediate the efficient adaptive immune response (Makarova et al., 2018). Upon providing aptly sized prespacers *in vitro*, Cas1-2 of *Streptococcus pyogenes* (type II-A) and *Enterococcus faecalis* (type-IIA) could swiftly integrate prespacers at the CRISPR locus (Wright and Doudna, 2016; Xiao et al., 2017). In contrast to this, spacer integration in Cas9 or Csn2 deprived *S. pyogenes* is extremely impaired (Heler et al., 2015; Wei et al., 2015b). Being an effector nuclease, Cas9 recognizes the PAM during target cleavage. Further, several studies have revealed that the Cas1-2 of *S. pyogenes* physically interacts with Cas9 and utilizes the PAM selection ability of Cas9 to identify potential prespacers (Heler et al., 2015; Ka et al., 2018). On the contrary, our experiments with *E. coli* (type I-E) CRISPR adaptation machinery suggest that the Cas1-2 complex alone is sufficient to recognize the PAM. Nuclease footprints of Cas1-2 on long DNA duplexes (63 bp) revealed that Cas1-2 intrinsically prefers PAM containing sequences during prespacer selection (Figure 3). Interestingly, mutation of PAM residues in longer DNA fragments leads to the non-specific capture of substrates at multiple points (Figure 3). This observation perhaps explains the reason for fractional uptake of spacers that are devoid of PAM in *E. coli* (Wt in Figure 4E). Uptake of prespacers with erroneous PAM preference is not uncommon among various type I systems (Datsenko et al., 2012; Li et al., 2017; Rao et al., 2017; Savitskaya et al., 2013; Shmakov et al., 2014). Such imprecision in spacer acquisition was shown to heighten the immune response by switching to primed adaptation (Datsenko et al., 2012; Jackson et al., 2019; Musharova et al., 2019; Musharova et al., 2018). A close inspection of the crystal structure of Cas1-2-prespacer complex highlights the features that could possibly lead to precise scaling and PAM selection of prespacers. A platform formed by the interaction of a Cas2 dimer with two Cas1 dimers on either side houses the 23 bp duplex region of prespacer (Nunez et al., 2015a; Wang et al., 2015). Stationed at either end of this duplex is the aromatic ring of Y22 residue that stacks the prespacer at the border of the Cas1 catalytic groove. Here, Y22 interaction acts as a wedge and directs the 3’-overhang to position its 5^th^ nt at the catalytic site. Besides positioning the PAM sequence at the active site of the integrase complex, Y22 guided meticulous placement of DNA substrate seems to dictate the length of prespacer (Nunez et al., 2015a; Wang et al., 2015). Furthermore, the flexible C-terminal tail of Cas1 is moulded around the PAM sequence of prespacer. The absence of such molecular architecture upon mutating the PAM region hints at the role of C-terminal tail in PAM recognition (Wang et al., 2015). Deployment of Cas1-2 variants that encompass either deletion of Cas1 C-terminal tail (ΔC) or alanine replacement of Cas1 Y22 in spacer integration assays, helped to unveil the role of these structural entities in determining the PAM selection (Figure 4D). As shown here, the deletion of C-terminal tail resulted in impaired PAM recognition (Figure 4A) and led to an uptake of prespacers that were lacking PAM (ΔC in Figure 4E). Moreover, a close inspection into the structures of Cas1 from *Archeoglobus fulgidus* (type I-A; PDB: 4N06) (Kim et al., 2013), *Pyrococcus horikoshii* (type I-B; PDB ID: 4WJ0), *B. halodurans* (type I-C, Cas1 modelled using I-TASSER (Zhang, 2008)) and *E. coli* (type I-E; PDB ID: 5DQZ) (Wang et al., 2015) revealed a striking contrast between Cas1 C-terminal tail of type I-E and other subtypes (Figure 5). Here, the Cas1 C-terminal tail of *A. fulgidus*, *P. horikoshii* and *B. halodurans* comprises of 12 amino acids (aa) and noticeably the C-terminal tail of *E. coli* Cas1 was of 31 aa in length (compare Figure 5A, 5B and 5C with 5D). These observations were further bolstered by detailed analysis of the length of the C-terminal tail of Cas1 multiple sequence alignments comprising of type I-A, I-B, I-C and I-E with reference to respective Cas1 structural features of *A. fulgidus*, *P. horikoshii*, *B. halodurans* and *E. coli* (average length of C-terminal tail in type I-E = 29 aa vs 12 aa in type I-A, I-B and I-C; Figure 5E). Coincidentally, CRISPR-Cas subtypes with shorter Cas1 C-terminal tails such as type I-A, I-B and I-C encompass Cas4 (Makarova et al., 2018) and previous studies suggest the indispensability of Cas4 in prompting PAM specificity (Kieper et al., 2018; Lee et al., 2018; Rollie et al., 2018; Shiimori et al., 2018). In contrast to these systems, it appears that the extended C-terminal tail of Cas1 in *E. coli* (type I-E) compensates for the lack of Cas4 by guiding the PAM detection. Regards to the variant Y22A, though the prespacer integration is unaffected *in vitro* (Figure S5B), in line with the previous obervations (Nunez et al., 2015a), a rampant decrease in spacer integration efficiency was observed *in vivo* (Figure 4C). Owing to the differences in affinity to P63 (Y22A K_D_ = 6.22 μM vs Wt K_D_ = 831.4 nM) (Figure S6D), Y22A conferred reduced protection against the action of nucleases (Figure 4A). In Wt, Y22 residue seems to offer a better grip on bound DNA against the nuclease action (Figure 4A). As Y22A could lack such interactions with substrates, nucleases seem to dislodge the bound DNA seamlessly (Figure 4A). This action appears to limit the substrate availability and impede the spacer integration *in vivo* (Figure 4C). Despite the reduction in spacer acquisition potential, Y22A showed high fidelity towards recognition of ‘5’-AAG-3’’ PAM (Y22A in Figure 4E). Owing to the absence of Cas1 C-terminal tail restructuring, prespacers lacking the PAM sequence are possibly held weakly by Cas1-2. In Y22A, the clutch on the substrate could be further enfeebled by the absence of tyrosine mediated stacking interactions. Due to these factors, cellular nucleases could deftly displace the PAM deficient prespacers than those with a PAM. Such bias displayed by Y22A in PAM selection appears to confer the improved fidelity for spacer integration.

**Figure 5.**
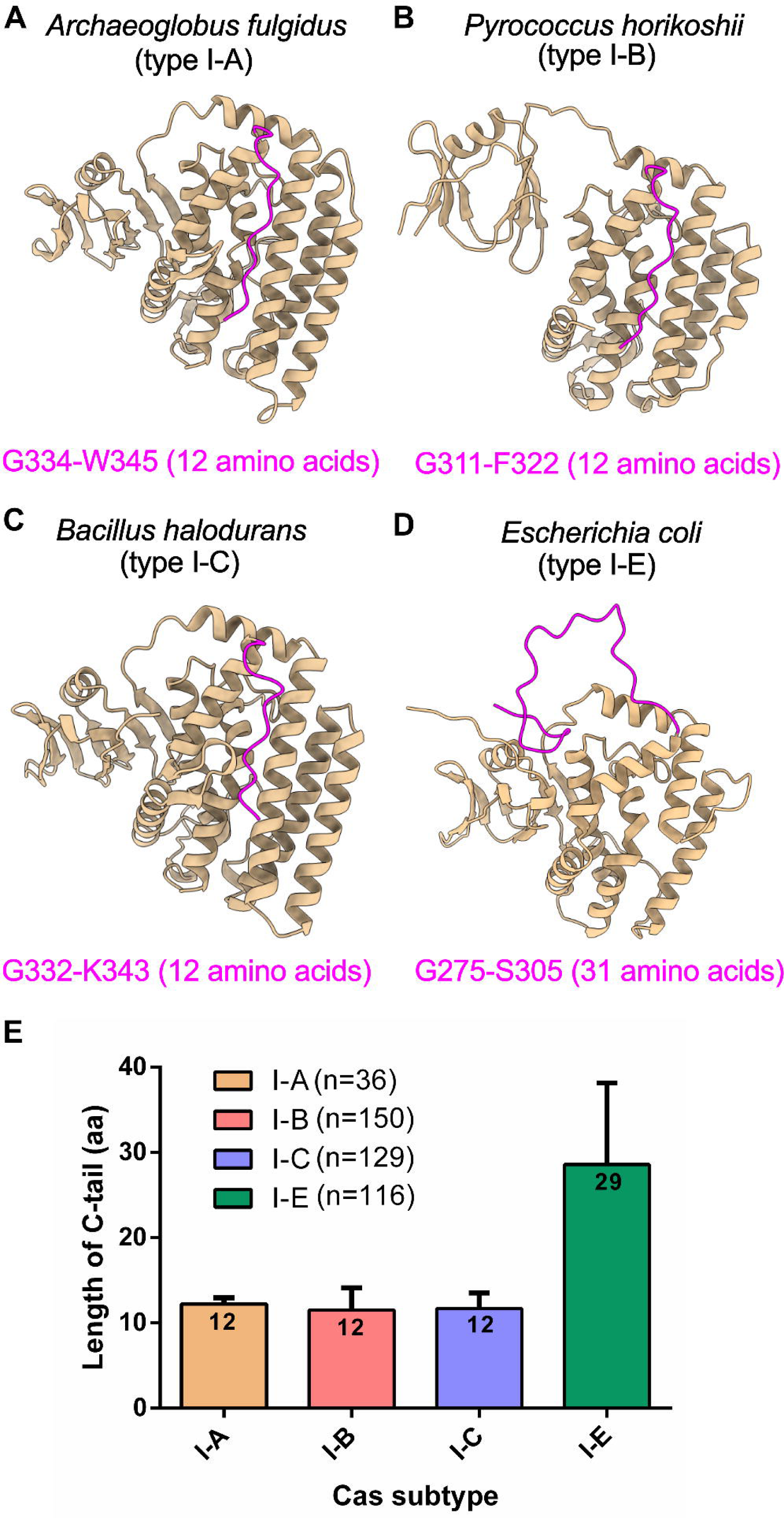
Type I-E Cas1 harbours extended C-terminal tail. (A-D) Structures highlighting Cas1 C-terminal tail (in magenta) of *Archeoglobus fulgidus* (type I-A; PDB ID: 4N06) (A), *Pyrococcus horikoshii* (type I-B; PDB ID: 4WJ0) (B), *Bacillus halodurans* (type I-C; predicted using homology based modelling tool I-TASSER; *vide.* method section) and *Escherichia coli* (type I-E; PDB ID: 5DQZ) (D) are portrayed. The amino acids corresponding to start and end position of C-terminal tail are displayed at the bottom of the respective structures. (E) Bar graph representing the length differences among Cas1 of I-A, I-B, I-C and I-E subtypes is displayed. Bars corresponding to each Cas subtype are shown in different colours and average length of amino acids in C-terminal tail of each subtype is indicated in the respective bars. ‘n’ corresponds to the number of Cas1 sequences from each subtype that are considered for the analysis (*vide* methods section).

Despite the differences in the PAM preference, the length of the acquired spacers corresponding to the Cas1-2 variants was predominantly observed to be 33 nt (Figure 4D) These findings negate the involvement of Y22 stacking interactions in deciding the prespacer boundary. Like in type V-C, where a structure of mini integrase complex constituted by Cas1 tetramer prefers short (18 bp) spacers (Wright et al., 2019), in *E. coli*, Cas1-2 structural framework alone appears to be a critical parameter in gauging the length of spacers (Nunez et al., 2015a; Wang et al., 2015).

Based on the state of the art and our current research work, we propose here an updated model for prespacer capture and integration in type I-E (Figure 6). Cas1-2 integrase captures the DNA fragments generated by nuclease activities of RecBCD and/or Cas3. Here, intrinsic specificity conferred by C-terminal tail localizes the Cas1-2 integrase at PAM regions. Upon interacting with PAM region, Cas1-2 hold protects the prespacer boundaries, whereas, the exposed ends of the DNA fragments were trimmed by the action of cellular nucleases. This Cas1-2-prespacer nucleoprotein complex loads at the binding site of CRISPR locus that is generated due to IHF mediated CRISPR leader remodelling. Upon recognizing the site of integration at the CRISPR locus, the nucleophic attacks by 3’-OH ends of prespacer (1^st^ attack at leader-repeat1 junction and 2^nd^ attack at repeat1-spacer1 junction) ligates itself as a new spacer (S0) (Figure 6). Overall, our study highlights the mechanism by which CRISPR adaptation machinery can manoeuvre itself to associate with host nucleases to achieve successful prespacer integration by offsetting the deficit of specialized prespacer trimming Cas4 nucleases.

**Figure 6.**
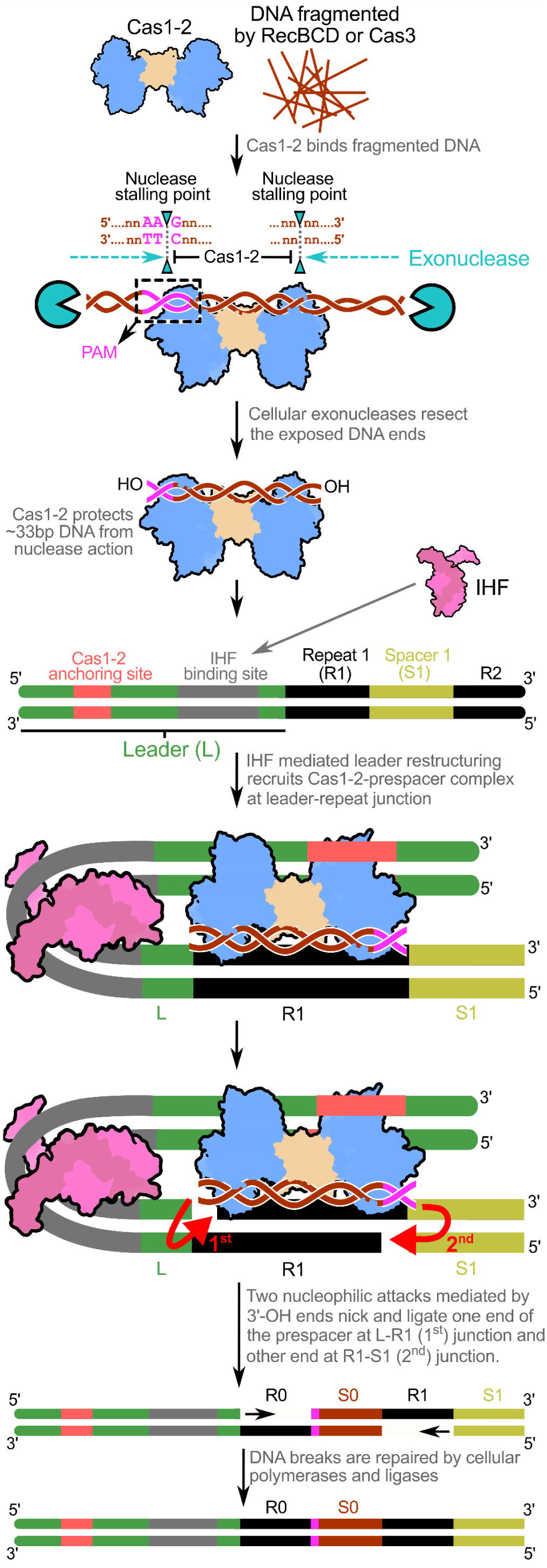
Model depicting the prespacer processing and integration in *E. coli*. Cas1-2 complex captures the DNA fragments (in brown) that arise from the nuclease activity of RecBCD or Cas3. Owing to its intrinsic specificity, Cas1-2 is recruited at PAM (5’-AAG-3’ in magenta) on the fragmented DNA. Upon this, cellular exonucleases (in cyan) degrade the exposed DNA ends, whereas Cas1-2 hold acts as a roadblock and stalls the exonuclease. This action protects 33 bp prespacer region starting with ‘G’ residue of the PAM sequence (5’-AAG-3’ in magenta). At the CRISPR locus, IHF (in pink) mediated deformation of IHF binding site (in grey) generates a cognate binding site for Cas1-2-prespacer complex by juxtaposing Cas1-2 anchoring site (in coral red) with leader(L)-repeat1(R1) junction. Consequent to localization of the Cas1-2-prespacer complex at the L-R1 junction, nucleophilic attack by 3’-OH ends prompt homing of prespacer by transesterification. Here, 1^st^ nucleophilic attack ligates the non-PAM end of the prespacer to 5’ end of the repeat at L-R1 junction in the top strand. On the other hand, 2^nd^ nucleophilic attack ligates ‘G’ residue derived from the PAM (in magenta) to 5’ end of the repeat at R1-spacer1(S1) junction in the bottom strand. Following this, cellular polymerases and ligases repair the DNA lesions to generate a CRISPR locus expanded with new spacer (S0) and duplicated repeat (R0).

## METHOD DETAILS

### Construction of plasmids

Lists of plasmids, strains and oligonucleotides used in this study are detailed in the supplementary tables 1-3.

Genes encoding IHFα, IHFβ, Cas1 and Cas2 were amplified using *Escherichia coli* K-12 MG1655 genomic DNA as template. To generate p1R-IHFαβ, a bicistronic cassette encoding IHFα and IHFβ was amplified and inserted at the SspI site of p1R. p13SR-Cas1 was generated by inserting an amplicon encoding Cas1 at SspI site of p13SR and pMS-Cas2 was created by introducing an amplicon encoding Cas2 between BamHI/HindIII sites of pMS (Guerrero et al., 2015), respectively. Bicistronic cassettes (*cas1-cas2*) expressing 6X Histidine tagged Wt and 5M Cas1 were amplified using pCas1-2[K] (Diez-Villasenor et al., 2013) and pMut89 (Shipman et al., 2016) as templates, respectively. Amplified fragments encoding Wt and 5M Cas1-2 were inserted between NcoI/NotI sites of pCas1-2[K] to generate pCas1-2H and p5M, respectively. PCR based mutagenesis was used to generate pY22A and pΔC that express Y22A and ΔC variants of Cas1, respectively. All constructs were verified by Sanger sequencing.

### Expression and purification of proteins

IHF and Cas1 were purified as described before (Yoganand et al., 2017). In order to purify Cas2, *E. coli* BL21(DE3) harbouring pMS-Cas2 was grown in auto-induction media supplemented with 100 µg/ml kanamycin at 37°C, 180 rpm. Upon reaching 0.6 OD_600_, temperature was shifted to 16°C and thereafter the growth and induction was continued for 16 hours. Subsequently, cells were harvested and washed 2X times with Buffer 1A (20 mM HEPES–NaOH pH 7.4, 500 mM KCl and 10% glycerol). Bacterial pellet was resuspended in Buffer 1A containing 1 mM PMSF and the cells were lysed by sonication. Here, Cas2 encompasses 6X Histidine tagged MBP-SUMO as an N-terminal fusion and a Strep-II tag on the C-terminal end. The clarified fraction of the lysate was applied to a 5 ml MBPTrap HP column (GE healthcare) and was followed by a washing step with Buffer 1A. Thereafter, the bound proteins were eluted with Buffer 1A containing 10 mM maltose. Eluted fractions were mixed with SUMO protease (Ulp1_403-621_) (in 400:1 ratio of His-MBP-SUMO-Cas2-strep: Ulp1_403-621_) (Guerrero et al., 2015) and the incubation was continued for 60 min at 25°C. Following this, the mixture was loaded onto 5 ml HiTrap IMAC HP column (GE Healthcare) 5X times to facilitate binding of Histidine tagged MBP-SUMO-Cas2-strep, MBP-SUMO and Ulp1_403-621_. Column flow-through containing Cas2-strep was concentrated using a centrifugal membrane filter (Sartorius). To relieve any trace protein contaminants, the concentrated sample was loaded onto HiLoad Superdex 200 pg gel filtration column (GE healthcare), that is pre-equilibrated with Buffer 1B (20 mM HEPES–NaOH pH 7.4, 150 mM KCl and 10% glycerol). Eluted fractions containing Cas2-strep were pooled, concentrated, snap frozen in liquid nitrogen and stored at −80°C until required.

Integrase complex comprising of untagged Cas1 and C-terminal 6X Histidine tagged Cas2 was expressed and purified as described before (Moch et al., 2017) with minor modifications. Here, *E. coli* BL21(DE3) transformed using pCas1-2H was grown in 2XYT broth supplemented with 100 µg/ml spectinomycin at 37°C, 180 rpm till 0.6 OD_600_. Thereafter, the protein expression was induced by addition of 0.7 mM IPTG and the growth was continued at 25°C for 24 hours. Simultaneously, cells were harvested and washed 2X times with Buffer 2A (20 mM HEPES–NaOH pH 7.4, 150 mM KCl, 10% glycerol and 30 mM imidazole). The pellet was resuspended in Buffer 2A containing 1 mM PMSF and cells were lysed by sonication. Thereafter, the lysate was clarified and loaded onto a 5 ml HiTrap IMAC HP column (GE Healthcare) and was followed by a washing step with Buffer 2A. A linear gradient of imidazole (0.03–0.5 M) in Buffer 2A was applied to elute the proteins that were bound to the column resin. The purified fractions that contain the complex of Cas1-2 were pooled and concentrated using a centrifugal membrane filter (Sartorius). To remove trace protein contaminants and un-complexed Cas2, the concentrate was further purified using HiLoad Superdex 200 pg gel filtration column (GE healthcare) that is pre-equilibrated with Buffer 2B (20 mM HEPES–NaOH pH 7.4, 150 mM KCl and 10% glycerol). Eluted fractions containing Cas1-2 integrase were pooled, concentrated, snap frozen in liquid nitrogen and stored at −80°C until required. Similar procedure was implemented to purify 5M, ΔC and Y22A Cas1 variants of Cas1-2 from the IPTG induced *E. coli* BL21(DE3) cells that harbours p5M, pΔC and pY22A, respectively.

### *In vitro* integration assay

CRISPR DNA substrate that encompasses 69 bp leader and two repeat-spacer units of CRISPR locus 2.1 of *E. coli* was amplified using pCSIR-T (Diez-Villasenor et al., 2013) as a template. To generate various prespacers (P33, P23[3’-5], P23[5’-5], P23[3’-10], P63 and P63mPAM), respective oligonucleotides (Supplementary table T3) were mixed in a buffer containing 10 mM Tris-Cl pH 8.5. These mixtures were heated to 95°C and gradually allowed to cool to room temperature in order to facilitate the formation of duplex and partial-duplex prespacers. Whereas, in case of P33ss, a 33nt long single stranded oligonucleotide was used as prespacer. The *in vitro* integrase assays were performed as previously described (Yoganand et al., 2017) with minor modifications. Briefly, 210 nM of Cas1 or Cas2 or Cas1-2 (Wt or ΔC or Y22A or 5M) was mixed with 550 nM of desired prespacer and incubated at room temperature for 5 minutes. To this mixture, 0.5 µM of IHF and 21 nM of CRISPR DNA substrate were supplemented and incubation was continued at 37°C for 60 min in integrase buffer (20 mM HEPES–NaOH pH 7.4, 25 mM KCl, 10 mM MgCl_2_ and 1 mM DTT). Subsequently, the reaction mixtures were treated with 1 mg/ml Proteinase-K and the incubation was continued for 30 min at 37°C. The samples were then electrophoresed on 8% native acrylamide gels in 1X TBE at 4°C. These gels were stained with EtBr and were photographed using gel documentation system (Bio-Rad).

### Electrophoretic mobility shift assays

The binding of Cas1-2 with various prespacers was monitored using electrophoretic mobility shift assays. Here, 175 nM of desired prespacers were incubated with increasing concentration of Wt or a mutant of Cas1-2 (0, 0.15, 0.2, 0.25, 0.3, 0.6, 0.8, 2, 3, 4, 8 and 12 µM) in prespacer binding buffer (20 mM HEPES–NaOH pH 7.4, 125 mM KCl, 10 mM MgCl_2_ and 1 mM DTT) for 30 min at 37°C. Thereafter, an aliquot of sample containing 175 nM of prespacer and 12 µM Cas1-2 was mixed with 1 mg/ml Proteinase-K and incubation was continued at 37°C for 15 min. Subsequently, all the samples were directly loaded on 8% native acrylamide gel and electrophoresed in 1X TBE at 4°C. These gels were stained with EtBr and photographed using gel documentation system. Bound fraction for each sample in the gel was estimated by quantifying the amount of DNA at each band using densitometric analysis (Bound fraction of prespacer (%) at X µM Cas1-2 = [(Amount of DNA in absence of Cas1-2 – Amount of unbound DNA at X µM Cas1-2] / [Amount of DNA in absence of Cas1-2)]*100). To estimate dissociation constants (K_D_) for each EMSA, the resulting plots of bound fraction (%) against Cas1-2 concentration was fitted to a non-linear equation: y = B_max_*x/(K_D_ + x) (where x, y, B_max_ and K_D_ represents Cas1-2 concentration (μM), bound fraction (%), maximum concentration of Cas1-2 bound to prespacer and dissociation constant, respectively).

### Exonuclease treatment of Cas1-2 bound DNA fragments

Exonuclease treatment was performed to identify the extent of protection conferred by binding of Cas1-2 on to a long DNA fragment. 40 µl of 0.5 µM P63 and 6 µM of either Cas1 or Cas2 or Cas1-2 in prespacer binding buffer were incubated at 37°C for 45 min. Subsequently, 20 µl aliquots of these samples were supplemented with a mixture containing 3 units each of T5 exonuclease (NEB) and Exonuclease III (NEB) and incubation was continued for 60 min at 37°C. Thereafter, all the samples were mixed with equal volume of denaturation buffer that contains 200 mM Tris–Cl pH 8.3, 200 mM Boric acid, 20 mM EDTA, 0.05 % SDS and 8 M Urea followed by heating at 95°C for 15 min. These samples were loaded on pre-heated 20% denaturation acrylamide gels that were maintained at 50°C and electrophoresed in 1X TBE. Subsequently, gels were stained with EtBr and visualized using gel documentation system

### Exonuclease footprinting

T5 exonuclease mediated footprinting was performed to identify the interaction boundaries of Cas1-2 on longer prespacer DNA fragments. Here, 40 µl of 0.5 µM of desired fluorescein labelled P63 variant (P63T*, P63B*, P63mPAMT* and P63mPAMB*) was mixed with 6 µM of Wt or one of the mutant variant of Cas1-2 (Y22A, ΔC and 5M) in prespacer binding buffer and incubated for 45 min at 37°C. Subsequently, 20 µl aliquots of these samples were supplemented with 3 units of T5 exonuclease and incubation was continued for 60 min at 37°C. Thereafter, all the samples were mixed with equal volume of denaturation buffer that contains 200 mM Tris–Cl pH 8.3, 200 mM Boric acid, 20 mM EDTA, 0.05 % SDS and 8 M Urea followed by heating at 95°C for 15 min. These samples were loaded on pre-heated 20% denaturation acrylamide gels that were maintained at 50°C and electrophoresed in 1X TBE. Subsequently, gels were directly visualized using gel documentation system.

### Spacer acquisition assays

The *in vivo* spacer acquisition assays were performed as described previously (Yoganand et al., 2017; Yosef et al., 2012) with minor modifications. *E. coli* IYB5101 transformed using plasmids (pCas1-2H or p5M or pY22A or pΔC) that express Wt or a mutant of Cas1-2 was subjected to three cycles of growth and induction in LB media supplemented with 100 μg/ml spectinomycin, 0.2% L-arabinose and 0.1 mM IPTG for 16 hours at 37°C. After each cycle, cultures were diluted to 1:300 times with fresh LB media containing aforementioned supplements and the growth was continued for 16 hours. Thereafter, genomic DNA was isolated according to manufacturer’s protocol (HiPurA bacterial genomic DNA purification kit, Himedia) and this was used as a template for PCR to monitor the spacer integration at CRISPR 2.1. All the PCR amplified samples were resolved on 1.5% agarose gels to identify the DNA bands corresponding to parental and expanded arrays (parental array + nx61 bp), where n is a positive integer. DNA quantities corresponding to parental and expanded array were quantified by densitometric analysis. Utilizing these values, percentage of spacer integration for each Cas1-2 variant was estimated (% integration = [(Amount of expanded array) / (Amount of parental array + Amount of expanded array)]*100).

### High-throughput sequencing and analysis

In order to understand the effect of Cas1-2 mutants on prespacer scaling and PAM selectivity, high-throughput sequencing was performed to derive the sequences of newly incorporated prespacers. Expanded CRISPR arrays corresponding to the expression of each Cas1-2 variant were extracted from the agarose gels (QIAquick Gel Extraction Kit, Qiagen). Approximately 200 ng of each PCR product was further purified using HighPrep magnetic beads (MAGBIO). These purified samples were subjected to DNA end repair and adaptor ligation using Illumina-compatible NEXTflex Rapid DNA sequencing kit (BIOO Scientific, Austin, Texas, U.S.A.). Subsequently, the ligated DNA products were purified with HighPrep magnetic beads and further enrichment was achieved by 8 cycles of PCR with Illumina-compatible primers (NEXTFlex DNA sequencing kit). These amplicons were subjected to an additional step of purification with HighPrep magnetic beads and were sequenced on a Miseq 300 paired end platform.

The paired-end reads were subjected to several pre-processing steps described as follows. Firstly, both F and R reads with Phred score less than 20 were removed by utilizing fastq_quality_trimmer from the FASTX-toolkit-version-0.0.13. The remaining F and R reads were trimmed in paired-end mode to remove F [5’-AGATCGGAAGAGCACACGTCTGAACTCCAGTCA-3’] and R [5’-AGATCGGAAGAGCGTCGTGTAGGGAAAGAGTGT-3’] adapter sequences using Cutadapt-1.18 (Martin, 2011). Following these, the leader proximal spacer sequence (S0) was selectively retrieved in FASTA format. These S0 sequences that were derived from *E. coli* expressing WT and mutants of Cas1-2 were searched against plasmid (pCas1-2[K]) (Diez-Villasenor et al., 2013) and *E. coli* K-12 MG1655 genome (GenBank assembly accession:GCA_000005845.2), respectively, using BLASTN (Johnson et al., 2008). From the BLAST hits, we identified the location of spacer sequences on plasmid and *E. coli* K-12 MG1655 genome, respectively and extracted the triplet sequence corresponding to PAM. The conservation of PAM was analysed using WEB-LOGO (Crooks et al., 2004). For all sequence manipulations including the extraction of S0 and PAM sequences, we employed custom written python codes utilising the Biopython library (Cock et al., 2009).

### Analysis of Cas1 C-terminal tail across CRISPR-Cas types I-A, I-B, I-C and I-E

Locus ID’s corresponding to Cas1 genes of type I-A, I-B, I-C and I-E were derived from the previous study (Makarova et al., 2015). Utilizing these identifiers we extracted Cas1 protein sequences of 36 type I-A, 150 type I-B, 129 type I-C and 116 type I-E organisms from Entrez database. Subsequent to this, we performed multiple sequence alignments in T-COFFEE web server (Notredame et al., 2000) for Cas1 proteins of each subtype separately. Utilizing Cas1 crystal structures of *A. fulgidus* (type I-A; PDB ID: 4N06), *P. horikoshii* (type I-B; PDB ID: 4WJ0) and *E. coli* (type I-E; PDB ID: 5DQZ) as a reference, C-terminal tail residues of each Cas1 was extracted from their multiple sequence alignments. Owing to the absence of type I-C Cas1 crystal structure in PDB database, Cas1 (BH0341) structure of *B. halodurans* was predicted using I-TASSER web server (Zhang, 2008). By using structures corresponding to PDB ID: 3LFX, 4N06 and 2YZS as threading templates, I-TASSER predicted five different models for *B. halodurans* Cas1. Among these, the model with highest confidence score (C-score = 1.25) was used as a structural reference for predicting the C-terminal tail residues from multiple sequence alignment of 129 type I-C Cas1 proteins.

## DATA AVAILABILITY

Reads obtained from high-throughput sequencing have been deposited in the Sequence Read Archive (SRA) under accession number: PRJNA527928.

## ACKNOWLEDGEMENTS

This work was supported in part by grants from Department of Biotechnology (DBT) [BT/PR15925/NER/95/141/2015 and BT/08/IYBA/2014/05] and Science and Engineering Research Board (SERB) [YSS/2014/000286]. *E. coli* IYB5101 strain was a kind gift from Dr. Udi Qimron, Tel Aviv University, Israel; Vectors pET StrepII TEV LIC cloning vector (p1R) (Addgene #29664) and pET StrepII TEV co-transformation cloning vector (p13SR) (Addgene #48328) were a kind gift from Dr. Scott Gradia, QB3 MacroLab, University of California, USA; pLJSRSF7 (pMS) (Addgene #64693) and pFGET19_Ulp1(Addgene #64697) were a kind gift from Dr. Hideo Iwai, University of Helsinki, Finland; pWUR 1+2 tetO mut89 (pMut89) (Addgene #80102) was a kind gift from Dr. George M. Church, Harvard University, USA; pCSIR-T and pCas1-2[K] were a kind gift from Dr. F.J.M. Mojica, University of Alicante, Spain. We sincerely acknowledge the gracious gesture of the aforementioned scientists for sharing their bacterial strains and plasmids. We also thank all the members of MAB lab for their suggestions at the bench work and critical comments on the manuscript.

## AUTHOR CONTRIBUTIONS

K.N.R.Y. and B.A conceived the research problem; all authors designed the assays; K.N.R.Y. and M.M. performed the experiments; B.A. analysed the high-throughput sequencing data; K.N.R.Y. and B.A. wrote the manuscript; B.A. arranged for funds and oversaw the project.

## REFERENCES

Al-Attar, S., Westra, E.R., van der Oost, J., and Brouns, S.J. (2011). Clustered regularly interspaced short palindromic repeats (CRISPRs): the hallmark of an ingenious antiviral defense mechanism in prokaryotes. Biol. Chem. 392, 277–289.

Almendros, C., Nobrega, F.L., McKenzie, R.E., and Brouns, S.J.J. (2019). Cas4–Cas1 fusions drive efficient PAM selection and control CRISPR adaptation. Nucleic Acids Res. https://doi.org/10.1093/nar/gkz217

Amitai, G., and Sorek, R. (2016). CRISPR-Cas adaptation: insights into the mechanism of action. Nat. Rev. Microbiol. 14, 67–76.

Arslan, Z., Hermanns, V., Wurm, R., Wagner, R., and Pul, U. (2014). Detection and characterization of spacer integration intermediates in type I-E CRISPR-Cas system. Nucleic Acids Res. 42, 7884–7893.

Barrangou, R., Fremaux, C., Deveau, H., Richards, M., Boyaval, P., Moineau, S., Romero, D.A., and Horvath, P. (2007). CRISPR provides acquired resistance against viruses in prokaryotes. Science 315, 1709–1712.

Cock, P.J.A., Antao, T., Chang, J.T., Chapman, B.A., Cox, C.J., Dalke, A., Friedberg, I., Hamelryck, T., Kauff, F., Wilczynski, B., et al. (2009). Biopython: freely available Python tools for computational molecular biology and bioinformatics. Bioinformatics 25, 1422–1423.

Crooks, G.E., Hon, G., Chandonia, J.M., and Brenner, S.E. (2004). WebLogo: a sequence logo generator. Genome Res. 14, 1188–1190.

Datsenko, K.A., Pougach, K., Tikhonov, A., Wanner, B.L., Severinov, K., and Semenova, E. (2012). Molecular memory of prior infections activates the CRISPR/Cas adaptive bacterial immunity system. Nat. Commun. 3, 945.

Diez-Villasenor, C., Guzman, N.M., Almendros, C., Garcia-Martinez, J., and Mojica, F.J. (2013). CRISPR-spacer integration reporter plasmids reveal distinct genuine acquisition specificities among CRISPR-Cas I-E variants of Escherichia coli. RNA Biol. 10, 792–802.

Dillingham, M.S., and Kowalczykowski, S.C. (2008). RecBCD enzyme and the repair of double-stranded DNA breaks. Microbiol. Mol. Biol. Rev. 72, 642–671.

Drabavicius, G., Sinkunas, T., Silanskas, A., Gasiunas, G., Venclovas, Č., and Siksnys, V. (2018). DnaQ exonuclease-like domain of Cas2 promotes spacer integration in a type I-E CRISPR-Cas system. EMBO Rep. 19, e45543.

Fagerlund, R.D., Wilkinson, M.E., Klykov, O., Barendregt, A., Pearce, F.G., Kieper, S.N., Maxwell, H.W.R., Capolupo, A., Heck, A.J.R., Krause, K.L., et al. (2017). Spacer capture and integration by a type I-F Cas1–Cas2-3 CRISPR adaptation complex. Proc. Natl. Acad. Sci. U. S. A. 114, E5122.

Fineran, P.C., and Charpentier, E. (2012). Memory of viral infections by CRISPR-Cas adaptive immune systems: acquisition of new information. Virology 434, 202–209.

Goren, M.G., Doron, S., Globus, R., Amitai, G., Sorek, R., and Qimron, U. (2016). Repeat Size Determination by Two Molecular Rulers in the Type I-E CRISPR Array. Cell Rep. 16, 2811–2818.

Guerrero, F., Ciragan, A., and Iwaï, H. (2015). Tandem SUMO fusion vectors for improving soluble protein expression and purification. Protein Expr. Purif. 116, 42–49.

Heler, R., Samai, P., Modell, J.W., Weiner, C., Goldberg, G.W., Bikard, D., and Marraffini, L.A. (2015). Cas9 specifies functional viral targets during CRISPR-Cas adaptation. Nature 519, 199–202.

Hille, F., Richter, H., Wong, S.P., Bratovič, M., Ressel, S., and Charpentier, E. (2018). The Biology of CRISPR-Cas: Backward and Forward. Cell 172, 1239–1259.

Hochstrasser, M.L., and Doudna, J.A. (2015). Cutting it close: CRISPR-associated endoribonuclease structure and function. Trends Biochem. Sci. 40, 58–66.

Horvath, P., and Barrangou, R. (2010). CRISPR/Cas, the immune system of bacteria and archaea. Science 327, 167–170.

Ivancic-Bace, I., Cass, S.D., Wearne, S.J., and Bolt, E.L. (2015). Different genome stability proteins underpin primed and naive adaptation in E. coli CRISPR-Cas immunity. Nucleic Acids Res. 43, 10821–10830.

Jackson, S.A., Birkholz, N., Malone, L.M., and Fineran, P.C. (2019). Imprecise Spacer Acquisition Generates CRISPR-Cas Immune Diversity through Primed Adaptation. Cell Host Microbe 25, 250–260.e254.

Jackson, S.A., McKenzie, R.E., Fagerlund, R.D., Kieper, S.N., Fineran, P.C., and Brouns, S.J. (2017). CRISPR-Cas: Adapting to change. Science 356, eaal5056.

Johnson, M., Zaretskaya, I., Raytselis, Y., Merezhuk, Y., McGinnis, S., and Madden, T.L. (2008). NCBI BLAST: a better web interface. Nucleic Acids Res. 36, W5–W9.

Ka, D., Jang, D.M., Han, B.W., and Bae, E. (2018). Molecular organization of the type II-A CRISPR adaptation module and its interaction with Cas9 via Csn2. Nucleic Acids Res. 46, 9805–9815.

Kieper, S.N., Almendros, C., Behler, J., McKenzie, R.E., Nobrega, F.L., Haagsma, A.C., Vink, J.N.A., Hess, W.R., and Brouns, S.J.J. (2018). Cas4 Facilitates PAM-Compatible Spacer Selection during CRISPR Adaptation. Cell Rep. 22, 3377–3384.

Kim, T.Y., Shin, M., Huynh Thi Yen, L., and Kim, J.S. (2013). Crystal structure of Cas1 from Archaeoglobus fulgidus and characterization of its nucleolytic activity. Biochem. Biophys. Res. Commun. 441, 720–725.

Kunin, V., Sorek, R., and Hugenholtz, P. (2007). Evolutionary conservation of sequence and secondary structures in CRISPR repeats. Genome Biol. 8, R61.

Künne, T., Kieper, S.N., Bannenberg, J.W., Vogel, A.I.M., Miellet, W.R., Klein, M., Depken, M., Suarez-Diez, M., and Brouns, S.J.J. (2016). Cas3-Derived Target DNA Degradation Fragments Fuel Primed CRISPR Adaptation. Mol. Cell 63, 852–864.

Lee, H., Zhou, Y., Taylor, D.W., and Sashital, D.G. (2018). Cas4-Dependent Prespacer Processing Ensures High-Fidelity Programming of CRISPR Arrays. Mol. Cell 70, 48–59.e45.

Levy, A., Goren, M.G., Yosef, I., Auster, O., Manor, M., Amitai, G., Edgar, R., Qimron, U., and Sorek, R. (2015). CRISPR adaptation biases explain preference for acquisition of foreign DNA. Nature 520, 505–510.

Li, M., Gong, L., Zhao, D., Zhou, J., and Xiang, H. (2017). The spacer size of I-B CRISPR is modulated by the terminal sequence of the protospacer. Nucleic Acids Res. 45, 4642–4654.

Li, M., Wang, R., and Xiang, H. (2014a). Haloarcula hispanica CRISPR authenticates PAM of a target sequence to prime discriminative adaptation. Nucleic Acids Res. 42, 7226–7235.

Li, M., Wang, R., Zhao, D., and Xiang, H. (2014b). Adaptation of the Haloarcula hispanica CRISPR-Cas system to a purified virus strictly requires a priming process. Nucleic Acids Res. 42, 2483–2492.

Liu, T., Liu, Z., Ye, Q., Pan, S., Wang, X., Li, Y., Peng, W., Liang, Y., She, Q., and Peng, N. (2017). Coupling transcriptional activation of CRISPR-Cas system and DNA repair genes by Csa3a in Sulfolobus islandicus. Nucleic Acids Res. 45, 8978–8992.

Makarova, K.S., Haft, D.H., Barrangou, R., Brouns, S.J., Charpentier, E., Horvath, P., Moineau, S., Mojica, F.J., Wolf, Y.I., Yakunin, A.F., et al. (2011). Evolution and classification of the CRISPR-Cas systems. Nat. Rev. Microbiol. 9, 467–477.

Makarova, K.S., Wolf, Y.I., Alkhnbashi, O.S., Costa, F., Shah, S.A., Saunders, S.J., Barrangou, R., Brouns, S.J., Charpentier, E., Haft, D.H., et al. (2015). An updated evolutionary classification of CRISPR-Cas systems. Nat. Rev. Microbiol. 13, 722–736.

Makarova, K.S., Wolf, Y.I., and Koonin, E.V. (2018). Classification and Nomenclature of CRISPR-Cas Systems: Where from Here? The CRISPR Journal 1, 325–336.

Marraffini, L.A. (2015). CRISPR-Cas immunity in prokaryotes. Nature 526, 55–61.

Marraffini, L.A., and Sontheimer, E.J. (2008). CRISPR interference limits horizontal gene transfer in staphylococci by targeting DNA. Science 322, 1843–1845.

Marraffini, L.A., and Sontheimer, E.J. (2010). Self versus non-self discrimination during CRISPR RNA-directed immunity. Nature 463, 568–571.

Martin, M. (2011). Cutadapt removes adapter sequences from high-throughput sequencing reads. EMBnet J. 17, 10–12.

McGinn, J., and Marraffini, L.A. (2016). CRISPR-Cas Systems Optimize Their Immune Response by Specifying the Site of Spacer Integration. Mol. Cell 64, 616–623.

McGinn, J., and Marraffini, L.A. (2019). Molecular mechanisms of CRISPR-Cas spacer acquisition. Nat. Rev. Microbiol. 17, 7–12.

Moch, C., Fromant, M., Blanquet, S., and Plateau, P. (2017). DNA binding specificities of Escherichia coli Cas1-Cas2 integrase drive its recruitment at the CRISPR locus. Nucleic Acids Res. 45, 2714–2723.

Mohr, G., Silas, S., Stamos, J.L., Makarova, K.S., Markham, L.M., Yao, J., Lucas-Elío, P., Sanchez-Amat, A., Fire, A.Z., Koonin, E.V., et al. (2018). A Reverse Transcriptase-Cas1 Fusion Protein Contains a Cas6 Domain Required for Both CRISPR RNA Biogenesis and RNA Spacer Acquisition. Mol. Cell 72, 700–714.e708.

Mojica, F.J., Diez-Villasenor, C., Garcia-Martinez, J., and Almendros, C. (2009). Short motif sequences determine the targets of the prokaryotic CRISPR defence system. Microbiology 155, 733–740.

Musharova, O., Klimuk, E., Datsenko, K.A., Metlitskaya, A., Logacheva, M., Semenova, E., Severinov, K., and Savitskaya, E. (2017). Spacer-length DNA intermediates are associated with Cas1 in cells undergoing primed CRISPR adaptation. Nucleic Acids Res. 45, 3297–3307.

Musharova, O., Sitnik, V., Vlot, M., Savitskaya, E., Datsenko, K.A., Krivoy, A., Fedorov, I., Semenova, E., Brouns, S.J.J., and Severinov, K. (2019). Systematic analysis of Type I-E Escherichia coli CRISPR-Cas PAM sequences ability to promote interference and primed adaptation. Mol. Microbiol. https://doi.org/10.1111/mmi.14237.

Musharova, O., Vyhovskyi, D., Medvedeva, S., Guzina, J., Zhitnyuk, Y., Djordjevic, M., Severinov, K., and Savitskaya, E. (2018). Avoidance of Trinucleotide Corresponding to Consensus Protospacer Adjacent Motif Controls the Efficiency of Prespacer Selection during Primed Adaptation. mBio 9, e02169–02118.

Notredame, C., Higgins, D.G., and Heringa, J. (2000). T-coffee: a novel method for fast and accurate multiple sequence alignment. J. Mol. Biol. 302, 205–217.

Nunez, J.K., Bai, L., Harrington, L.B., Hinder, T.L., and Doudna, J.A. (2016). CRISPR Immunological Memory Requires a Host Factor for Specificity. Mol. Cell 62, 824–833.

Nunez, J.K., Harrington, L.B., Kranzusch, P.J., Engelman, A.N., and Doudna, J.A. (2015a). Foreign DNA capture during CRISPR-Cas adaptive immunity. Nature 527, 535–538.

Nuñez, J.K., Kranzusch, P.J., Noeske, J., Wright, A.V., Davies, C.W., and Doudna, J.A. (2014). Cas1–Cas2 complex formation mediates spacer acquisition during CRISPR–Cas adaptive immunity. Nat. Struct. Mol. Biol. 21, 528–534.

Nunez, J.K., Lee, A.S., Engelman, A., and Doudna, J.A. (2015b). Integrase-mediated spacer acquisition during CRISPR-Cas adaptive immunity. Nature 519, 193–198.

Pul, U., Wurm, R., Arslan, Z., Geissen, R., Hofmann, N., and Wagner, R. (2010). Identification and characterization of E. coli CRISPR-cas promoters and their silencing by H-NS. Mol. Microbiol. 75, 1495–1512.

Punetha, A., Yoganand, K.N.R., Nimkar, S., and Anand, B. (2018). Cutting it right: Plasticity and strategy of crispr RNA specific nucleases. Proc. Indian Natl. Sci. Acad. (B Biol. Sci.) 84, 455–477.

Radovcic, M., Killelea, T., Savitskaya, E., Wettstein, L., Bolt, E.L., and Ivancic-Bace, I. (2018). CRISPR-Cas adaptation in Escherichia coli requires RecBCD helicase but not nuclease activity, is independent of homologous recombination, and is antagonized by 5’ ssDNA exonucleases. Nucleic Acids Res. 46, 10173–10183.

Rao, C., Chin, D., and Ensminger, A.W. (2017). Priming in a permissive type I-C CRISPR-Cas system reveals distinct dynamics of spacer acquisition and loss. RNA (New York, N.Y.) 23, 1525–1538.

Rollie, C., Graham, S., Rouillon, C., and White, M.F. (2018). Prespacer processing and specific integration in a Type I-A CRISPR system. Nucleic Acids Res. 46, 1007–1020.

Rollie, C., Schneider, S., Brinkmann, A.S., Bolt, E.L., and White, M.F. (2015). Intrinsic sequence specificity of the Cas1 integrase directs new spacer acquisition. Elife 4, e08716.

Savitskaya, E., Semenova, E., Dedkov, V., Metlitskaya, A., and Severinov, K. (2013). High-throughput analysis of type I-E CRISPR/Cas spacer acquisition in E. coli. RNA Biol. 10, 716–725.

Semenova, E., Savitskaya, E., Musharova, O., Strotskaya, A., Vorontsova, D., Datsenko, K.A., Logacheva, M.D., and Severinov, K. (2016). Highly efficient primed spacer acquisition from targets destroyed by the Escherichia coli type I-E CRISPR-Cas interfering complex. Proc. Natl. Acad. Sci. U. S. A. 113, 7626–7631.

Shiimori, M., Garrett, S.C., Graveley, B.R., and Terns, M.P. (2018). Cas4 Nucleases Define the PAM, Length, and Orientation of DNA Fragments Integrated at CRISPR Loci. Mol. Cell 70, 814–824.e816.

Shipman, S.L., Nivala, J., Macklis, J.D., and Church, G.M. (2016). Molecular recordings by directed CRISPR spacer acquisition. Science 353, aaf1175.

Shmakov, S., Savitskaya, E., Semenova, E., Logacheva, M.D., Datsenko, K.A., and Severinov, K. (2014). Pervasive generation of oppositely oriented spacers during CRISPR adaptation. Nucleic Acids Res. 42, 5907–5916.

Silas, S., Mohr, G., Sidote, D.J., Markham, L.M., Sanchez-Amat, A., Bhaya, D., Lambowitz, A.M., and Fire, A.Z. (2016). Direct CRISPR spacer acquisition from RNA by a natural reverse transcriptase-Cas1 fusion protein. Science 351, aad4234.

Sinkunas, T., Gasiunas, G., Fremaux, C., Barrangou, R., Horvath, P., and Siksnys, V. (2011). Cas3 is a single-stranded DNA nuclease and ATP-dependent helicase in the CRISPR/Cas immune system. EMBO J. 30, 1335–1342.

Sternberg, S.H., Richter, H., Charpentier, E., and Qimron, U. (2016). Adaptation in CRISPR-Cas Systems. Mol. Cell 61, 797–808.

Swarts, D.C., Mosterd, C., van Passel, M.W., and Brouns, S.J. (2012). CRISPR interference directs strand specific spacer acquisition. PLoS One 7, e35888.

Wang, J., Li, J., Zhao, H., Sheng, G., Wang, M., Yin, M., and Wang, Y. (2015). Structural and Mechanistic Basis of PAM-Dependent Spacer Acquisition in CRISPR-Cas Systems. Cell 163, 840–853.

Wang, R., Li, M., Gong, L., Hu, S., and Xiang, H. (2016). DNA motifs determining the accuracy of repeat duplication during CRISPR adaptation in Haloarcula hispanica. Nucleic Acids Res. 44, 4266–4277.

Wei, Y., Chesne, M.T., Terns, R.M., and Terns, M.P. (2015a). Sequences spanning the leader-repeat junction mediate CRISPR adaptation to phage in Streptococcus thermophilus. Nucleic Acids Res. 43, 1749–1758.

Wei, Y., Terns, R.M., and Terns, M.P. (2015b). Cas9 function and host genome sampling in Type II-A CRISPR-Cas adaptation. Genes Dev. 29, 356–361.

Westra, E.R., Pul, U., Heidrich, N., Jore, M.M., Lundgren, M., Stratmann, T., Wurm, R., Raine, A., Mescher, M., Van Heereveld, L., et al. (2010). H-NS-mediated repression of CRISPR-based immunity in Escherichia coli K12 can be relieved by the transcription activator LeuO. Mol. Microbiol. 77, 1380–1393.

Wright, A.V., and Doudna, J.A. (2016). Protecting genome integrity during CRISPR immune adaptation. Nat. Struct. Mol. Biol. 23, 876–883.

Wright, A.V., Liu, J.J., Knott, G.J., Doxzen, K.W., Nogales, E., and Doudna, J.A. (2017). Structures of the CRISPR genome integration complex. Science 357, 1113–1118.

Wright, A.V., Wang, J.Y., Burstein, D., Harrington, L.B., Paez-Espino, D., Kyrpides, N.C., Iavarone, A.T., Banfield, J.F., and Doudna, J.A. (2019). A Functional Mini-Integrase in a Two-Protein-type V-C CRISPR System. Mol. Cell 73, 727–737.e723.

Xiao, Y., Ng, S., Nam, K.H., and Ke, A. (2017). How type II CRISPR-Cas establish immunity through Cas1-Cas2-mediated spacer integration. Nature 550, 137–141.

Yoganand, K.N.R., Sivathanu, R., Nimkar, S., and Anand, B. (2017). Asymmetric positioning of Cas1–2 complex and Integration Host Factor induced DNA bending guide the unidirectional homing of protospacer in CRISPR-Cas type I-E system. Nucleic Acids Res. 45, 367–381.

Yosef, I., Goren, M.G., and Qimron, U. (2012). Proteins and DNA elements essential for the CRISPR adaptation process in Escherichia coli. Nucleic Acids Res. 40, 5569–5576.

Yosef, I., Shitrit, D., Goren, M.G., Burstein, D., Pupko, T., and Qimron, U. (2013). DNA motifs determining the efficiency of adaptation into the Escherichia coli CRISPR array. Proc. Natl. Acad. Sci. U. S. A. 110, 14396–14401.

Zetsche, B., Gootenberg, J.S., Abudayyeh, O.O., Slaymaker, I.M., Makarova, K.S., Essletzbichler, P., Volz, S.E., Joung, J., van der Oost, J., Regev, A., et al. (2015). Cpf1 is a single RNA-guided endonuclease of a class 2 CRISPR-Cas system. Cell 163, 759–771.

Zhang, Y. (2008). I-TASSER server for protein 3D structure prediction. BMC Bioinformatics 9, 40.

Zhang, Z., Pan, S., Liu, T., Li, Y., and Peng, N. (2019). Cas4 nucleases can effect specific integration of CRISPR spacers. J. Bacteriol. https://doi.org/10.1128/JB.00747-18 JB.00747-00718.

