## Supplementary Information for "Fidelity of Prespacer Capture and Processing is Governed by the PAM Mediated Interaction of Cas1-2 Adaptation complex in *Escherichia coli*"

### Supplementary Figures

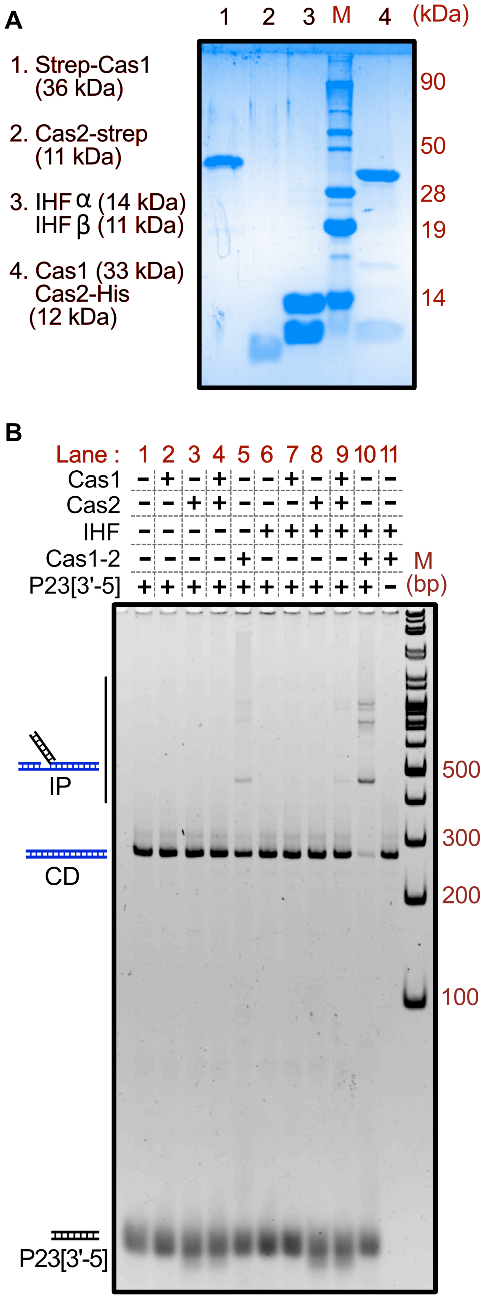


**Figure S1. Cas1-2 complex formation necessitates the integration of prespacers into CRISPR DNA.**

(A) 17% SDS-PAGE displays the purified Cas1, Cas2, IHF and Cas1-2. Molecular weight (in kDa) corresponding to proteins in each lane is shown on the left. The molecular weight marker (M) positions are shown on the right.

(B) Native gel displaying the proteinase K treated samples of spacer integration assay is shown. Absence (–) or presence (+) of Cas1, Cas2, IHF, Cas1-2 and prespacer P23[3'-5] is indicated on top of each lane. Positions of bands corresponding to CRISPR DNA (CD), integration product (IP) and prespacer (P23[3'-5]) are pictorially represented. The DNA molecular weight marker (M) positions are shown on the right. Integrated products were observed in the samples that contain Cas1-2 complex. Notably, supplementation of IHF greatly enhanced the homing of prespacers into the CRISPR DNA (compare lanes 5 and 9). Intriguingly, when Cas1, Cas2 and IHF were added individually, we observed the appearance of integration products albeit with reduced potential in comparison to the samples that contain IHF and Cas1-2 complex (compare lanes 9 and 10).

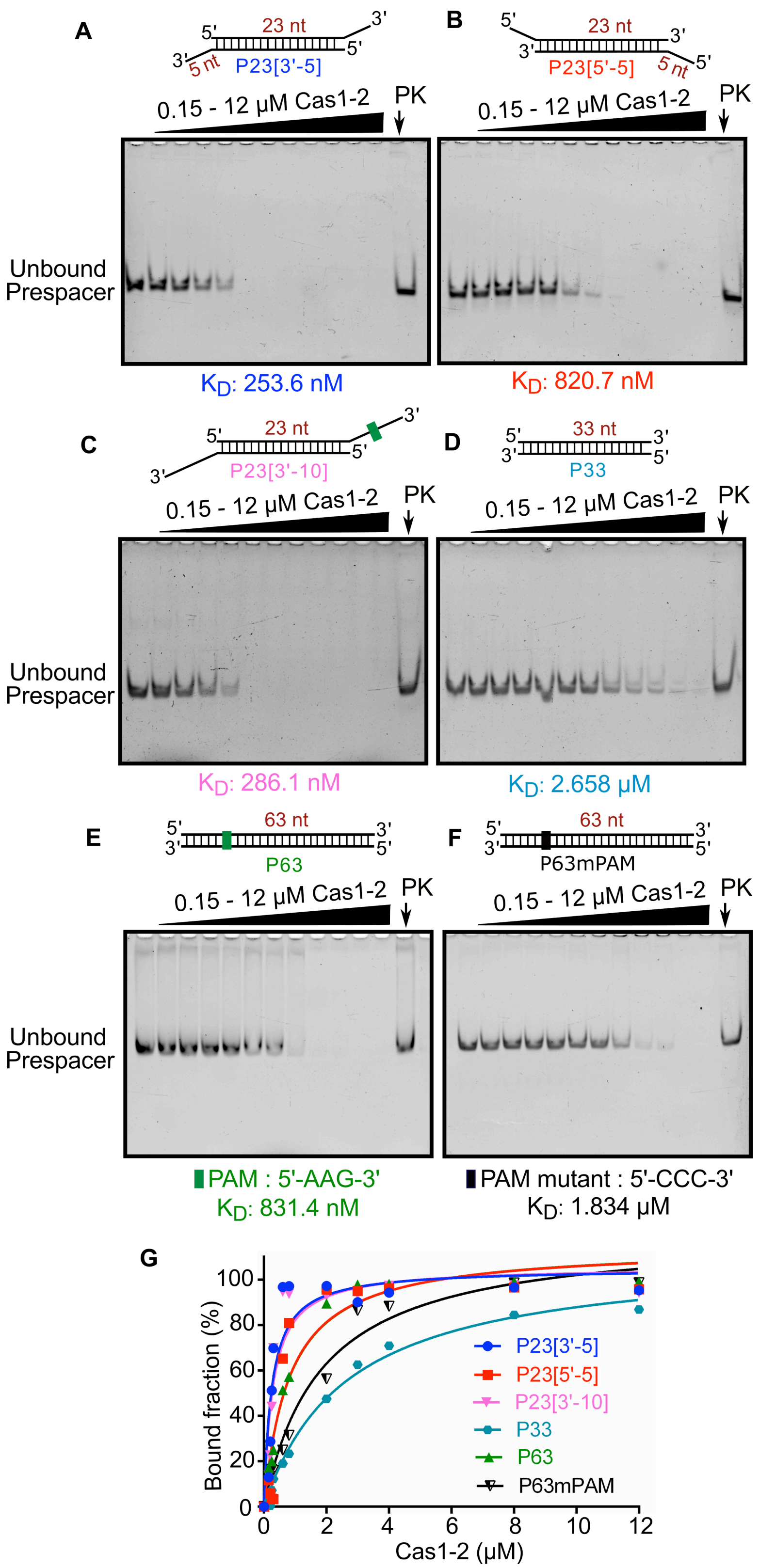


**Figure S2. Cas1-2 interacts with prespacers of varied lengths.**

(A-F) Native gels depicting the interactions of Cas1-2 with prespacers P23[3'-5] (A), P23[5'-5] (B), P23[3'-10] (C), P33 (D), P63 (E) and P63mPAM (F) are displayed. Schematic representation of the prespacer and the position of unbound prespacers are shown at the respective gel images. 100 nM of each prespacer DNA was incubated with increasing concentrations of Cas1-2 (0, 0.15, 0.2, 0.25, 0.3, 0.6, 0.8, 2, 3, 4, 8 and 12 μM). Because of the large size of Cas1-2/DNA complex, it wasn't entering the gel and therefore, we couldn't directly observe the shift in the mobility. Thus, an aliquot of the sample containing the mixture of 100 nM prespacer and 12 μM Cas1-2 was treated with proteinase K. Release of intact prespacer upon the proteinase K mediated digestion (lane PK in A-F) suggests the formation of Cas1-2-prespacer nucleoprotein complex.

(G) Plot of the bound fraction of prespacer (%) against Cas1-2 concentration (μM) for the electrophoretic mobility shift assays (EMSA in A-F) is displayed. The estimated $K_D$ values from the binding experiments are depicted at the bottom of the respective gels in (A-F).

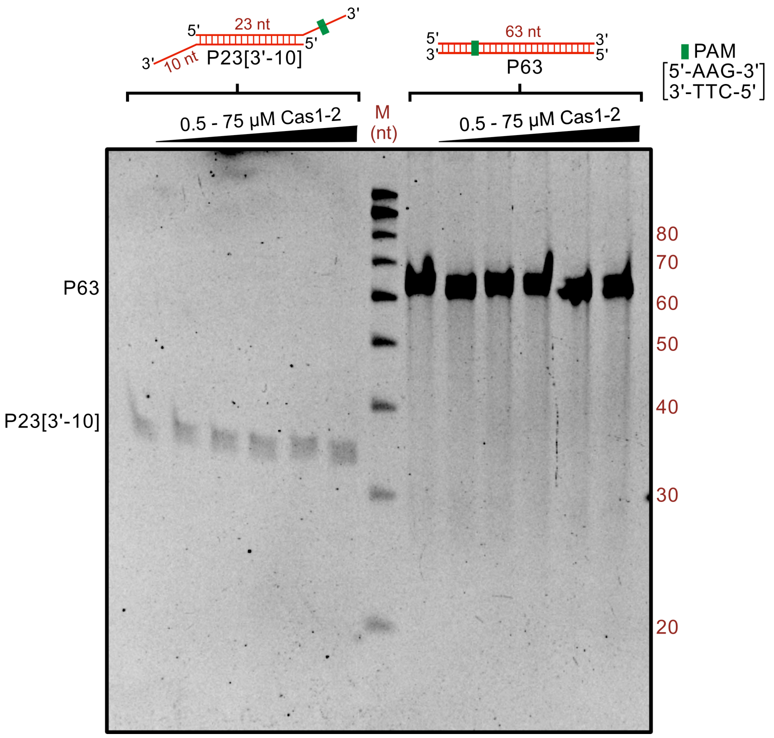


**Figure S3. Cas1-2 alone could not process the prespacers.** Denaturation gel depicting the interaction of 0.5 μM of prespacer DNA (P23[3'-10] and P63) with increasing concentrations of Cas1-2 (0, 0.5, 5, 10, 25 and 75 μM) is displayed. Pictorial depiction of P23[3'-10] and P63 is displayed on top of the respective lanes. Positions corresponding to each of the prespacer DNA and oligo marker (M) are shown on the sides of the gel. Though P23[3'-10] and P63 were capable of binding to Cas1-2 (Figure S2), generation of smaller DNA fragments was not observed with increasing concentrations of Cas1-2. These observations highlight that Cas1-2 by itself is inept in processing the prespacers.

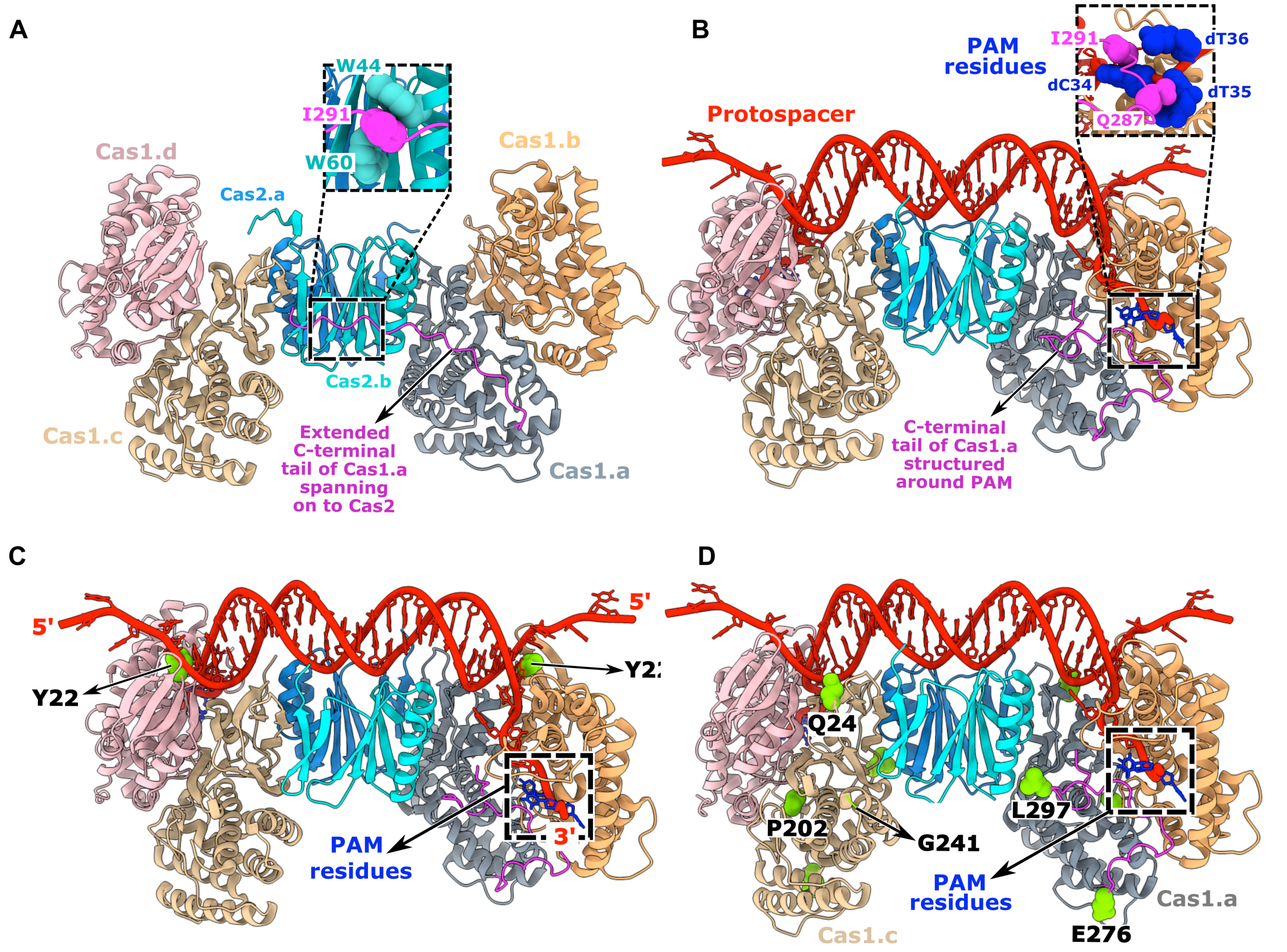


**Figure S4. Salient structural features of Cas1-2 determine the prespacer selection.**

(A-B) Structural comparison between apo-Cas1-2 integrase (PDB ID: 4P6I) (A) and Cas1-2-prespacer complex (PDB: 5DQZ) (B) is displayed. Four protomers of Cas1 (indicated as Cas1.a, Cas1.b, etc.) and two protomers of Cas2 (indicated as Cas2.a and Cas2.b) are shown in different colours. In apoCas1-2, an extended C-terminal tail of Cas1.a (G275-S305 in magenta) spans across Cas2. Here, I291 (Magenta spheres in close-up view at the top (A)) of the C-terminal tail is held by sandwiching between the Cas2 tryptophans (W44 and W60 as cyan spheres). Upon binding with prespacer, this disordered tail wraps around the PAM region at the prespacer border (B). Noticeably, Q287 and I291 residues of the C-terminal tail (magenta spheres in the close-up view) make a base specific contact with the PAM residues Cytosine 34 (dC34) and Thymidine 35 (dT35), respectively (Navy blue spheres in close-up view).

(C) Cas1-2-prespacer structure (PDB ID: 5DQZ) highlighting the interactions of Cas1 Y22 (as green spheres) with prespacer DNA (in red) is displayed. Here, the Y22 residues are positioned on either end of the Cas1-2 platform. These residues stack the borders of 23 bp prespacer duplex and ensure fork bifurcation that guides the PAM (in navy blue) containing 3’ overhang into the catalytic groove of Cas1-2 integrase.

(D) Amino acid residues corresponding to Cas1 mutations (Q24H, P202Q, G241D, E276D and L297Q) in 5M variant are displayed (green spheres) as part of the Cas1.a and Cas1.c protomers of Cas1-2-prespacer complex (PDB ID: 5DQZ). For clarity, residues corresponding to E276 and L297 of Cas1.a and Q24, P202 and G241 of Cas1.c are shown at their respective positions.

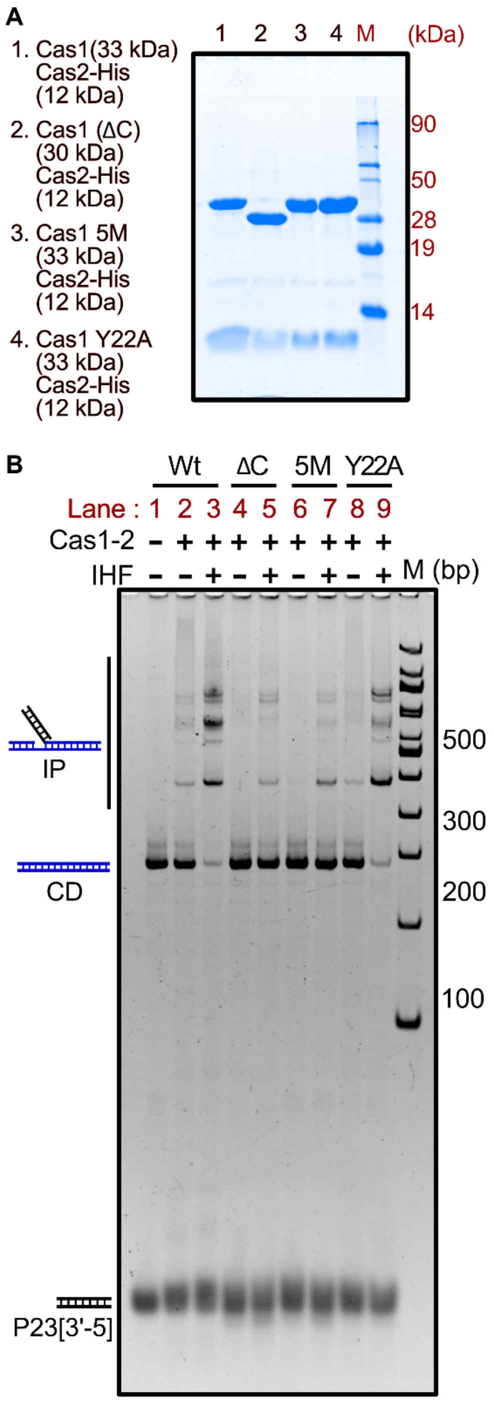


**Figure S5. Wt, ΔC, 5M and Y22A display prespacer integration into CRISPR DNA**

(A) 17% SDS-PAGE displaying purified Wt, ΔC, 5M and Y22A is shown. Molecular weight (kDa) corresponding to proteins in each lane is represented on the left. Protein molecular weight marker (M) positions are shown on the right.

(B) Native gel displaying the proteinase K treated samples of spacer integration assay is shown. Absence (–) or presence (+) of Cas1-2 and IHF in integration reaction is indicated on top of the respective lane. Positions of bands corresponding to CRISPR DNA (CD), integration product (IP) and prespacer (P23[3'-5]) are pictorially represented. DNA molecular weight marker (M) positions are shown on the right side. The appearance of slow migrating DNA (IP) in the samples containing IHF and Cas1-2 variant is indicative of integration activity displayed by each variant (Lanes 3, 5, 7 and 9).

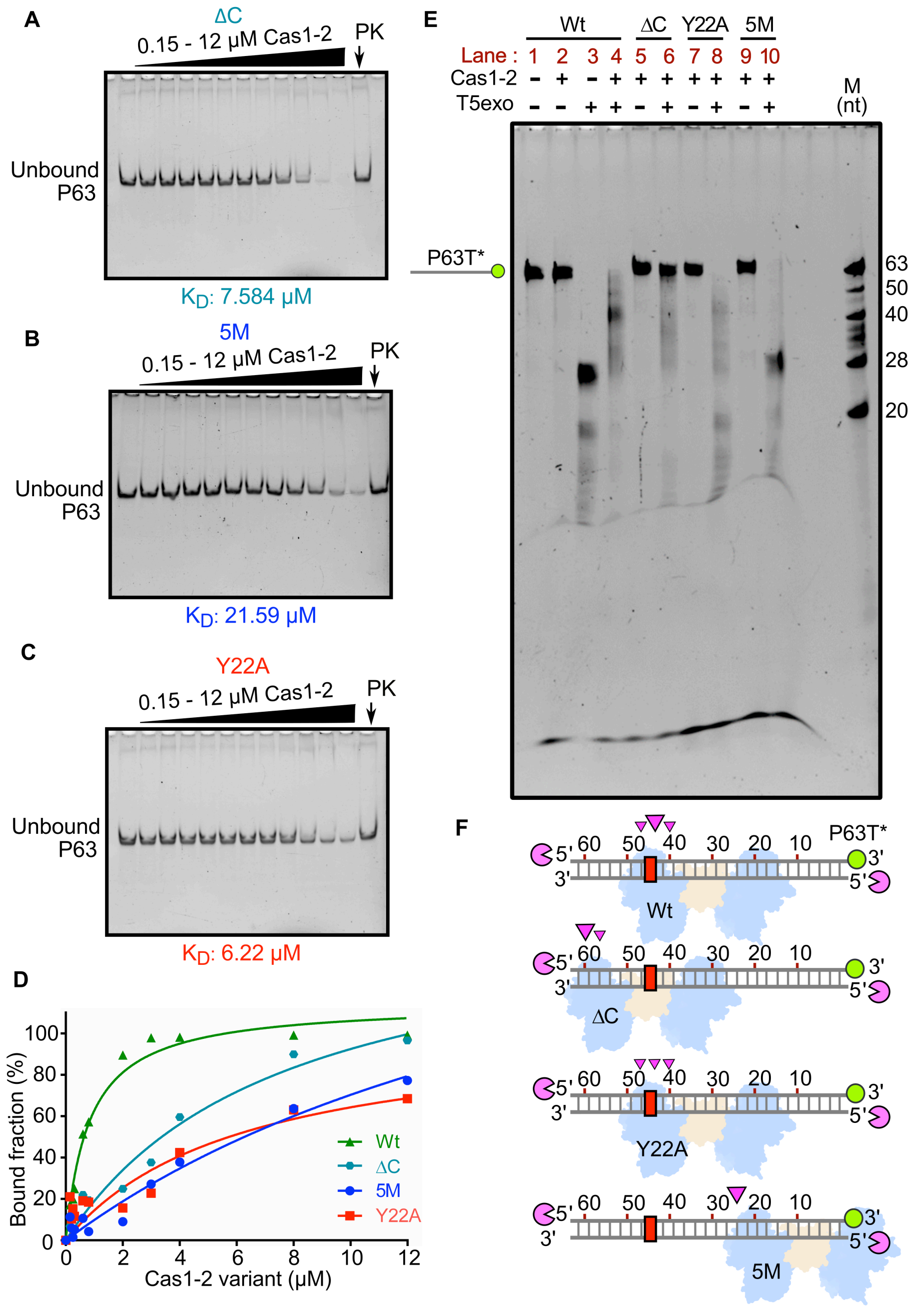


**Figure S6. Cas1-2 variants display differing specificities towards prespacers.**

(A-C) Native gels depicting the interactions of ΔC (A), 5M (B) and Y22A (C) with prespacer P63 are displayed. The variant of Cas1-2 employed and the position of unbound prespacers are shown at the respective gel images. 100 nM of P63 DNA was incubated with increasing concentrations of Cas1-2 (0, 0.15, 0.2, 0.25, 0.3, 0.6, 0.8, 2, 3, 4, 8 and 12 μM). Because of the large size of Cas1-2/DNA complex, it wasn't entering the gel and therefore, we couldn't directly observe the shift in the mobility. Thus, an aliquot of sample containing the mixture of 100 nM P63 and 12 μM of each Cas1-2 variant was treated with proteinase K. Release of intact prespacer upon the proteinase K mediated digestion (lane PK in A-C) suggests the formation of Cas1-2-prespacer complex.

(D) Plot of the bound fraction of prespacer (%) against Cas1-2 concentration (μM) for the electrophoretic mobility shift assays (EMSA in A-C) is displayed. The estimated $K_D$ values from the binding experiments are depicted at the bottom of the respective gels in (A-C). Whereas the data corresponding to bound fraction for Wt were obtained from Figure S2E.

(E) Denaturation gel depicting the T5exo treatment of Cas1-2 (Wt (lanes 1-4) or ΔC (lanes 5-6) or Y22A (lanes 7-8) or 5M (lanes 9-10)) bound fluorescein labelled P63T* is displayed. Presence (+) or absence (-) of each reaction component is labelled on top of each lane. Position of P63T* labelled DNA fragment is shown on the left, whereas oligo marker (M) positions are indicated on the right.

(F) Schematic illustration depicting the footprinting assay performed in (E) is displayed. DNA substrate P63T* (grey ladder), positions of 3' fluorescein label (green circle) and PAM region (red rectangle) are pictorially represented. Numbering on the DNA represents the distance (in nt) of particular position from the labelled end. T5 exo (magenta pie) is positioned at susceptible 5'-ends of DNA substrate. Positions of T5exo stalling points (magenta triangles) and binding sites of each variant of Cas1-2 (Wt or ΔC or

Y22A or 5M in blue and brown blobs) that are estimated from nuclease footprinting assay performed in (E) are pictorially indicated.

#### Supplementary Tables

**Supplementary table 1: Bacterial strains used in the study**

| *E. coli* strain | Genotype | Source |
|---|---|---|
| IYB5101 | F$^{-}$Δ(*araD-araB*)*567* Δ*lacZ4787* (::rrnB-3) λ$^{-}$ rph-1 Δ(*rhaD-rhaB*)*568 hsdR514 araB*::T7-RNAp-*tetA*, tet$^{r}$ | (Yosef et al., 2012) |
| DH5α | F$^{-}$ Φ80*lacZ*ΔM15 Δ*(lacZYA-argF) U169 recA1 endA1 hsdR17*($r_k^-$, $m_k^+$) *phoA supE44 thi-1 gyrA96 relA1 λ*$^{-}$ | Invitrogen |
| TOP10 | F$^{-}$ *mcrA* Δ(*mrr-hsdRMS-mcrBC*) ϕ80*lacZ*ΔM15 Δ *lacX74 recA1 araD139* Δ(*araleu*)7697 *galU galK rpsL* (str$^{r}$) *endA1 nupG* | Invitrogen |
| BL21(DE3) | F$^{-}$ *fhuA2 [lon] ompT gal (λ DE3) [dcm] ∆hsdS λ DE3 = λ sBamHIo ∆EcoRI-B int::(lacI::PlacUV5::T7 gene1) i21 ∆nin5* | NEB |

**Supplementary table 2: Plasmids used in the study**

| Plasmid name | Description | source |
|---|---|---|
| p1R | T7-lac inducible, ColE1 origin plasmid for expressing genes with N-terminally fused Strep-II tag. | Addgene #29664 |
| p13SR | T7-lac inducible, CloDF13 origin plasmid for expressing genes with N-terminally fused Strep-II tag. | Addgene #48328 |
| pMS | T7-lac inducible, RSF origin plasmid for expressing N-terminally 6X His-MBP-SUMO tagged genes. | Addgene #64693 |
| pFGET19_Ulp1 | Lac inducible plasmid for expression of 6X His tagged SUMO protease catalytic domain (Ulp1$_{403-621}$). | Addgene #64697 |
| pMut89 | CloDF13 origin plasmid for expressing Wt (Cas1-2) under T7-lac inducible promoter and 5M (Cas1(Q24H, P202Q, G241D, E276D, L297Q)-Cas2) under Tetracycline Inducible promoter. | Addgene #80102 |
| pCSIR-T | Plasmid containing CRISPR 2.1 array with 2 repeats | (Diez-Villasenor et al., 2013) |
| pCas1-2[K] | Plasmid expressing Cas1 and Cas2 of *E. coli* K-12 MG1655 | (Diez-Villasenor et al., 2013) |
| p1R-IHFαβ | Plasmid expressing N-terminally Strep-II tagged IHFα and untagged IHFβ under T7-lac promoter. | (Yoganand et al., 2017) |
| p13SR-Cas1 | Plasmid expressing N-terminally Strep-II tagged Cas1 under T7-lac promoter. | (Yoganand et al., 2017) |
| pMS-Cas2 | Plasmid expressing N-terminally 6X His-MBP-SUMO tagged, C-terminally Strep-II tagged Cas2 under T7-lac promoter. | This study |
| pCas1-2H | Plasmid expressing Cas1 (Wt), C-terminally 6X His tagged Cas2 under T7-lac promoter. | This study |
| p5M | Plasmid expressing Cas1 (5M: Q24H, P202Q, G241D, E276D, L297Q), C-terminally 6X His tagged Cas2 under T7-lac promoter. | This study |
| pY22A | Plasmid expressing Cas1 (Y22A), C-terminally 6X His tagged Cas2 under T7-lac promoter. | This study |
| pΔC | Plasmid expressing Cas1 (ΔC: ΔP279-S305), C-terminally 6X His tagged Cas2 under T7-lac promoter. | This study |

| Supplementary table 3: Oligonucleotides used in this study | | |
|---|---|---|
| **Oligo name** | **Sequence in 5’-3’ orientation** | **Purpose** |
| Ihfα 1R F | TACTTCCAATCCAATGCAATGGCGCTTACAAAAGCTGAAATGT | Generation of cassette encoding IHF with flanking sequences of SspI digested p1R (Yoganand et al., 2017). |
| Ihfα-RBS R | TTGGTCATGGTATATCTCCTTCTTAAAGTTAATTACTCGTCTTTGGGCGAAGC | |
| RBS-Ihfβ F | ATTAACTTTAAGAAGGAGATATACCATGACCAAGTCAGAATTGATAGAAAGACT | |
| Ihfβ 1R R | TTATCCACTTCCAATGTTATTATTAACCGTAAATATTGGCGCGATCGC | |
| Cas1 13SR F | TACTTCCAATCCAATGCAATGACCTGGCTTCCCCTTAATCC | Generation of DNA encoding Cas1 with flanking sequences of SspI digested p13SR (Yoganand et al., 2017). |
| Cas1 13SR R | TTATCCACTTCCAATGTTATTATCAGCTACTCCGATGGCCTGC | |
| Cas2 MS F | TCACAGAGAACAGATTGGTGGATCCGGAGGTATGAGTATGTTGGTCGTGGTCACTG | Generation of DNA encoding Cas2 with flanking sequences of BamHI and HindIII digested pMS. |
| Cas2 strep R | TTATTATTTTTCGAACTGCGGGTGGCTCCAAGCGCTAACAGGTAAAAAAGACACCAACCTTAAAC | |
| MS strep R | CTTTACCAGACTCGAGTGCGGCCGCAAGCTTTTATTATTTTTCGAACTGCGGGTGGC | |
| CDF wt Cas1 F | AACTTTAATAAGGAGATATACCATGGCCTGGCTTCCCCTTAATCCC | Generation of DNA constructs encoding Cas1-2 variants (Wt, 5M, ΔC, Y22A) with flanking sequences of NcoI and NotI digested pCas1-2[K]. |
| CDF Cas2 His R | TTATTAGTGATGGTGATGGTGATGAGCGCTAACAGGTAAAAAAGACACCAACCTTAAACC | |
| CDF His R | TTTCTTTACCAGACTCGAGTGCGGCCGCTTATTAGTGATGGTGATGGTGATGAGC | |
| ΔC Cas1-2 R | TTCAGGTGGGGCCGGCGGCTATTATTGTATTTCTCCAGCGGCAAGC | |
| ΔC Cas1-2 F | TAATAGCCGCCGGCCCCACCTGAA | |
| Cas1 Y22A F | TCGCGTCTCCATGATCTTTCTGCAAGCTGGGCAGATCGAT | |
| 2.1 array F | GGAAATGTTACATTAAGGTTGGTGGGTTG | Monitoring the spacer incorporation into the 2.1 CRISPR array during *in vivo* integration assay. |
| 2.1 array R | CGCTCAGAAATTCCAGACCCGATCC | |
| Leader F | TGCATCTCGAGGCATGCCTGCAGCGGCCGCCAGTGTGATGGATATCTG | Generation of CRISPR DNA substrates for *in vitro* integration assay. |
| Repeat2 R | GGATCGGAATTCGAGCTCGGTACCCACCTTTGGCTTCGGCTGC | |
| P23[3’-5] F | ATTTACTACTCGTTCTGGTGTTTCTCGT | These oligos were annealed to prepare P23[3’-5] prespacer. Whereas, P23[3’-5] F oligo alone is used as P33[ss] prespacer. |
| P23[3’-5] R | AAACACCAGAACGAGTAGTAAATTGGGC | |
| P23[3’-10] F | ATTTACTACTCGTTCTGGTGTTTCTCGTCAGGG | These oligos were annealed to prepare P23[3’-10] prespacer. |
| P23[3’-10] R | AAACACCAGAACGAGTAGTAAATTGGGCTTGAG | |
| P33 F | GCCCAATTTACTACTCGTTCTGGTGTTTCTCGT | These oligos were annealed to prepare |

| | | |
|---|---|---|
| P33 R | ACGAGAAACACCAGAACGAGTAGTAAATTGGGC | P33 prespacer. |
| P23[5'5] F | GCCCAATTTACTACTCGTTCTGGTGTTT | These oligos were annealed to prepare P23[5'-5] prespacer. |
| P23[5'5] R | ACGAGAAACACCAGAACGAGTAGTAAAT | |
| P63 F | CTCCGCGCTGTAGAAGTCACCATTGTTGTGCACGACGACATC ATTCCGTGGCGTTATCCAGCT | These oligos were annealed to prepare P63 prespacer. Residues corresponding to the PAM are depicted in green. |
| P63 R | AGCTGGATAACGCCACGGAATGATGTCGTCGTGCACAACAATG GTGACTTCTACAGCGCGGAG | |
| P63mPAM F | CTCCGCGCTGTAG**CCC**TCACCA**C**TGTTGTGCACGACGACA**C** CA**G**TCCGTGGCGTTATCCAGCT | These oligos were annealed to prepare P63mPAM prespacer. Mutated residues in the oligo are represented in bold. |
| P63mPAM R | AGCTGGATAACGCCACGGA**C**TG**G**TGTCGTCGTGCACAACA**G**T GGTGA**GGG**CTACAGCGCGGAG | |
| P63 3'FAM-F | CTCCGCGCTGTAGAAGTCACCATTGTTGTGCACGACGACATC ATTCCGTGGCGTTATCCAGCT - **FAM(3')** | This oligo was annealed with P63 R to generate P63 T*. |
| P63 3'FAM-R | AGCTGGATAACGCCACGGAATGATGTCGTCGTGCACAACAATG GTGACTTCTACAGCGCGGAG - **FAM(3')** | This oligo was annealed with P63 F to generate P63 B*. |
| P63mPAM 3'FAM F | CTCCGCGCTGTAG**CCC**TCACCA**C**TGTTGTGCACGACGACA**C** CA**G**TCCGTGGCGTTATCCAGCT - **FAM(3')** | This oligo was annealed with P63mPAM R to generate P63mPAM T*. |
| P63mPAM 3'FAM R | AGCTGGATAACGCCACGGA**C**TG**G**TGTCGTCGTGCACAACA**G**T GGTGA**GGG**CTACAGCGCGGAG - **FAM(3')** | This oligo was annealed with P63mPAM F to generate P63mPAM B*. |

#### References


Diez-Villasenor, C., Guzman, N.M., Almendros, C., Garcia-Martinez, J., and Mojica, F.J. (2013). CRISPR-spacer integration reporter plasmids reveal distinct genuine acquisition specificities among CRISPR-Cas I-E variants of Escherichia coli. RNA Biol. *10*, 792-802.

Yoganand, K.N.R., Sivathanu, R., Nimkar, S., and Anand, B. (2017). Asymmetric positioning of Cas1–2 complex and Integration Host Factor induced DNA bending guide the unidirectional homing of protospacer in CRISPR-Cas type I-E system. Nucleic Acids Res. *45*, 367-381.

Yosef, I., Goren, M.G., and Qimron, U. (2012). Proteins and DNA elements essential for the CRISPR adaptation process in Escherichia coli. Nucleic Acids Res. *40*, 5569-5576.
